# Multivalent Scaffolding by XIST E-Repeats Tunes RNA Binding Protein Cooperativity and Lowers the Phase Separation Threshold

**DOI:** 10.64898/2026.10.25.761723

**Authors:** Bryan Eusse, Alfredo Vidal-Ceballos, Andrew Meyer, R. Alexander Martino, Emma Bose, Karmella Haynes, Alisha N Jones

## Abstract

The long-noncoding RNA XIST coordinates the random silencing of one X-chromosome in female placental mammals as a class of chromosomal dosage compensation across sexes. Critical to this process is the diffusion of XIST foci across the 155 Mb of the X-chromosome to achieve 96% repression. The 3’ E-Repeat domain on XIST primarily drives this process through the recruitment of several RNA-binding proteins, which seed condensate formation for controlled distal spreading. To date, there is a lack of biophysical characterization of interactions between the E-Repeats and its protein binding partners. Herein, we define the interaction, *in vitro*, between E-Repeat RNA and proteins PTBP1, ELAVL1, TDP43, and hnRNPU by applying a combination of bioinformatics, *in silico* characterization, and biochemical assays. We designed minimal E-Repeat RNA constructs to study the molecular mechanisms of RBP recognition and the E-Repeat ensemble. We find that all proteins sustain nanomolar interactions with single E-Repeat RNA. Through a systematic domain study, we deduce protein domain-specific RNA binding affinities, providing insight into mechanisms of E-Repeat binding. With four E-Repeats, we find that PTBP1 and ELAVL1 form higher-order assemblies and demonstrate protein competition and co-occupancy, reflecting previous *in cellulo* CLIP experiments. Finally, we find that PTBP1 and ELAVL1 phase separate with the four-repeat minimal constructs under minimal *in vitro* conditions, recapitulating the cellular behavior crucial to XIST function. All together, our findings provide mechanistic insight into the amplified impact of XIST, which at approximately 1 molecule per megabase can modulate long-range chromatin organization, illuminating another aspect of lncRNA potency, and potentially informing engineered derivatives of XIST and other lncRNAs as genetic therapeutics.

## Introduction

The function of RNA has been linked to both its structural dynamics and its interactions with other biomacromolecules, including DNA, RNA, and protein. These functional attributes enable biological phenomena that include biomolecular condensate formation, precise subcellular localization, and the orchestration of complex transcriptional machinery^1,2^. Ultimately, these mechanisms support stabilized states that allow cells to maintain homeostasis. A classic paradigm of these structure-function relationships is the X-inactive specific transcript (XIST), a long noncoding RNA (lncRNA) that has been extensively studied to better understand how it functions to stably silence the inactive X-chromosome in placental females^3–5^. This transcript possesses multiple repetitive domains, A, B, C, D, E, and F, which interact with over 80 RNA-binding proteins (RBPs) to function as a transcriptional regulator^5,6^. The most well-characterized of these domains include the A, B, and E repeat domains (**Figure 1A**).

**Figure 1.**
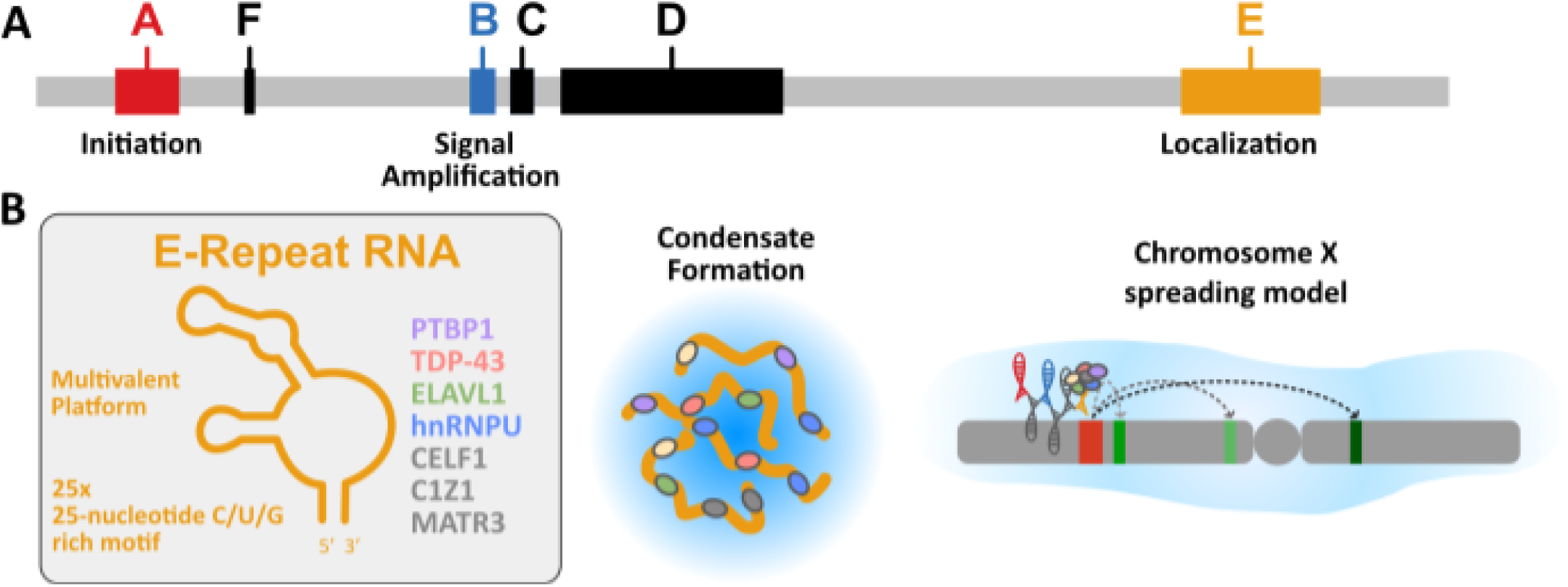
**A)** XIST’s multifunctionality is compartmentalized through its various repetitive domains. The E-Repeats are primarily responsible for the spreading of the XIST complex across the entire X-chromosome. **B)** The E-Repeats along with exon 7 act as a multivalent landing pad for various RBPs which seed phase separation and dispersal of the XIST cloud.

The A and B-Repeat domains are located at the 5’ end of XIST, and are responsible for silencing initiation and signal amplification, respectively. The A-Repeat domain adopts a structure that is conformationally dynamic; this domain directly interacts with numerous proteins, including SHARP and RBM15^7–9^. SHARP binds with nanomolar affinity to the A-repeats, which in turn recruits HDAC3 to deacetylate H3K27, a common repressive motif found on chromatin^10^. RBM15 binds the A-Repeats as a part of the METTL3, METTL14, WTAP (WMM) complex, to facilitate the methylation of adenosines (N6-methyladenosine) along the length of the transcript ^7,11,12^. Notably, in the absence of the A-Repeats (and its cognate RBPs), XIST diffusion still occurs, but transcriptional repression is abolished^13–15^. The B-Repeat domain is cytosine-rich in nature and is predicted to adopt several stem-loop motifs^16^. These C-rich patches facilitate a nanomolar affinity with hnRNPK via its C-terminal RGG domain^16^. The B-Repeat - hnRNPK interaction is critical for signal amplification, acting as a crucial scaffold that bridges XIST localization with the recruitment of polycomb repressive complexes 1 and 2 (PRC1 & PRC2)^17–19^. Upon binding to the B-Repeat scaffold, hnRNPK recruits PRC1, leading to the H2AK119 mono-ubiquinitation modification, which in turn recruits PRC2 to promote H3K27 trimethylation. Deletion of the B-repeats nullifies this signaling cascade and severely hampers the progression of heterochromatization *in vivo*.

The E-Repeats (∼1,500 nucleotides), the focus of this study, are located at the 3’ end of XIST and are responsible for the localization and spreading of XIST to chromatin (**Figure 1B**)^5,20,21^. The 2D structure of this domain has been chemically probed to reveal that, unlike the A and B-Repeats, the E-Repeats are largely unstructured, with stem loop domains interspersed amongst the single-stranded regions (**Figure 2A**)^22^. This is supported by the high chemical probing reactivities observed along the length of the RNA; high Shannon entropies suggest that the E-Repeats are conformationally dynamic, with a lack of defined structure, though the full structural ensemble remains unresolved. Crosslinking and chemical probing experiments have revealed

that the single-stranded regions of the E-Repeats overlap with binding sites for many proteins, including ELAVL1, PTBP1, TDP43, and hnRNPU^21–25^. These proteins engage with RNA through RNA recognition motifs (RRM) and RGG domains. For example, ELAVL1 (also known as HuR) possesses three RRMs; it binds AU-rich elements found in mRNA 3’ UTRs^26^. PTBP1 consists of four RRMs; RRMs 1 and 2 independently bind short, structured, pyrimidine-rich sequences^27^. This is in contrast to RRMs 3 and 4, which bind longer, single-stranded pyrimidine tracts^28,29^. TDP43 (also known as TARDBP) has two N-terminal RRMs that recognize and bind GU-rich mRNA intronic sequences. Unlike ELAVL1, PTBP1, TDP43, and hnRNPU (also known as SAF-A) possesses a disordered C-terminal RGG motif which binds GU-rich RNA^30–32^. This protein also contains an N-terminal DNA-binding SAP domain and internal SPRY and P-loop domains; the SAP domain is hypothesized to interact with chromatin, while the RGG domain binds XIST, potentially tethering them together^33^. Unlike the A and B repeats, the binding stoichiometry and overall assembly of the RNA-binding proteins contributing to XIST localization and spreading is largely uncharacterized.

In addition to harboring RNA-binding domains, ELAVL1, PTBP1, TDP43, and hnRNPU contain intrinsically disordered regions (IDRs), which are capable of facilitating liquid-liquid phase separation (LLPS) (**Figure 1B**)^34–36^. Some of these proteins can undergo simple coacervation in the absence of RNA (e.g. RBM15), whereas others have been found only to undergo phase separation in the presence of other bio-macromolecules (e.g., SHARP, TDP43 with hnRNPA1) and RNA, forming a complex coacervate. These disordered domains exist amongst the linker residues between the RRMs in ELAVL1 and PTBP1. HnRNPU has several disordered regions, including the RGG motif, which has previously been reported to promote aggregation of this RBP^37^. LLPS supports E-Repeat localization and the subsequent transcriptional silencing of the inactive X-chromosome^33,38,39^.

While previous studies focused on cellular knockouts, endogenous crosslinking, and RNA structural probing experiments to characterize these E-Repeat binding partners, such *in vivo* approaches are limited in their ability to decouple the intrinsic binding mechanisms of the individual RBPs from the context of DNA sequences, chromatin components, and the broader cellular environment. Indeed, an *in vitro* study with full-length PTBP1 and the E-Repeats revealed a near 200 nM binding affinity; *in vitro* phase separation assays revealed an RNA concentration dependence on condensate formation^38,40^. Isolating nucleoprotein complex dynamics outside the cellular environment laid the foundation for understanding how LLPS drives XIST-facilitated transcriptional silencing. This establishes the need for further, more in-depth studies investigating the structural and molecular properties that underlie the XIST E-Repeat-RBP interactions.

In this work, we build upon previous insights to dissect and characterize the interactions of PTBP1, ELAVL1, hnRNPU, and TDP43 with E-Repeat RNA by combining chemical probing deconvolution algorithms and biochemical and biophysical assays. Our work reveals distinct binding and interaction mechanisms for each cognate RBP, both in the dispersed and condensed phases. These insights advance our understanding of XIST localization, implicate E-Repeat-mediated protein scaffolding as an intrinsic, chromatin-agnostic property that might be ported to other contexts for cell engineering^41^, and further establishes XIST as a compelling therapeutic target for disease^14,42,43^.

## Results

### Structural analyses reveal a loosely structured, but dynamic XIST E-Repeat structure

To characterize the structural ensemble of the XIST E-Repeat domain, we performed deconvolution analysis on existing chemical probing data (**Supplemental Table 1**). DaVinci deconvolution analysis of chemical probing data across the ∼1,500-nucleotide domain identified three structural clusters in both *ex vivo* and in cellulo conditions (**Figure 2B**). *In cellulo*, the clusters were populated at 90.7, 9.3%, and <1%. Under *ex vivo* conditions, the three structural clusters were populated at 88.5%, 11.4%, and <1 % (**Figure 2B**). Across both conditions, the majority of the 25 E-Repeats remain unstructured, flanked by hairpins at the 5’ and 3’ extremities of the domain.

To determine how the molecular features of this ensemble govern RBP recognition, we synthesized minimal E-Repeat constructs containing one, two, three, or four tandem repeats (designated ERep1, ERep2, ERep3, and ERep4). The representative repeat sequence (5’-ggUUUCUUCCUUGCUUUGUGUGUCUA-3’) was adapted from an in-cell study (**Figure 2A**), and resembles the U-rich 5’-GUCUAUUUCUU-3’ motif previously identified as an RBP binding site for PTBP1 and CELF1^21,22^. We evaluated the secondary structure and conformational stability of these constructs using in silico folding algorithms and circular dichroism (CD) spectroscopy.

Folding of ERep1 predicted a thermodynamically unstable structure containing at most two base pairs among ten 5’ nucleotides, leaving the remaining sequence single-stranded. Increasing repeat length in ERep2, ERep3, and ERep4 did not increase secondary structure rigidity (**Figure 2C**). Boltzmann ensemble analysis similarly confirmed an absence of stable secondary structures; this was found to be true even when non-canonical base pairs were included within the data set to account for the high GU content of these RNAs^44^. Furthermore, biomolecular folding predictions using RNAduplex failed to identify complementary regions capable of forming duplexes between reads.

We validated these computational predictions experimentally using CD thermal denaturating assays (**Figure 2D**). Maximum intensity wavelengths selected for temperature ramps were 267 nm (ERep1), 270 nm (ERep2), 270 nm (ERep3), and 278 nm (ERep4). Thermal denaturation profiles for all four constructs exhibited high linearity (R^2^ ≥ 0.948), indicating an absence of cooperative unstacking or structured melting (**Figure 2D**). First derivative analysis revealed a broad, low intensity transition between 25°C and 30°C across all constructs, reflecting an ensemble of transient, weak secondary interactions (such as isolated GU base pairs) rather than stable tertiary folds (**Figure 2C, D**).

**Figure 2.**
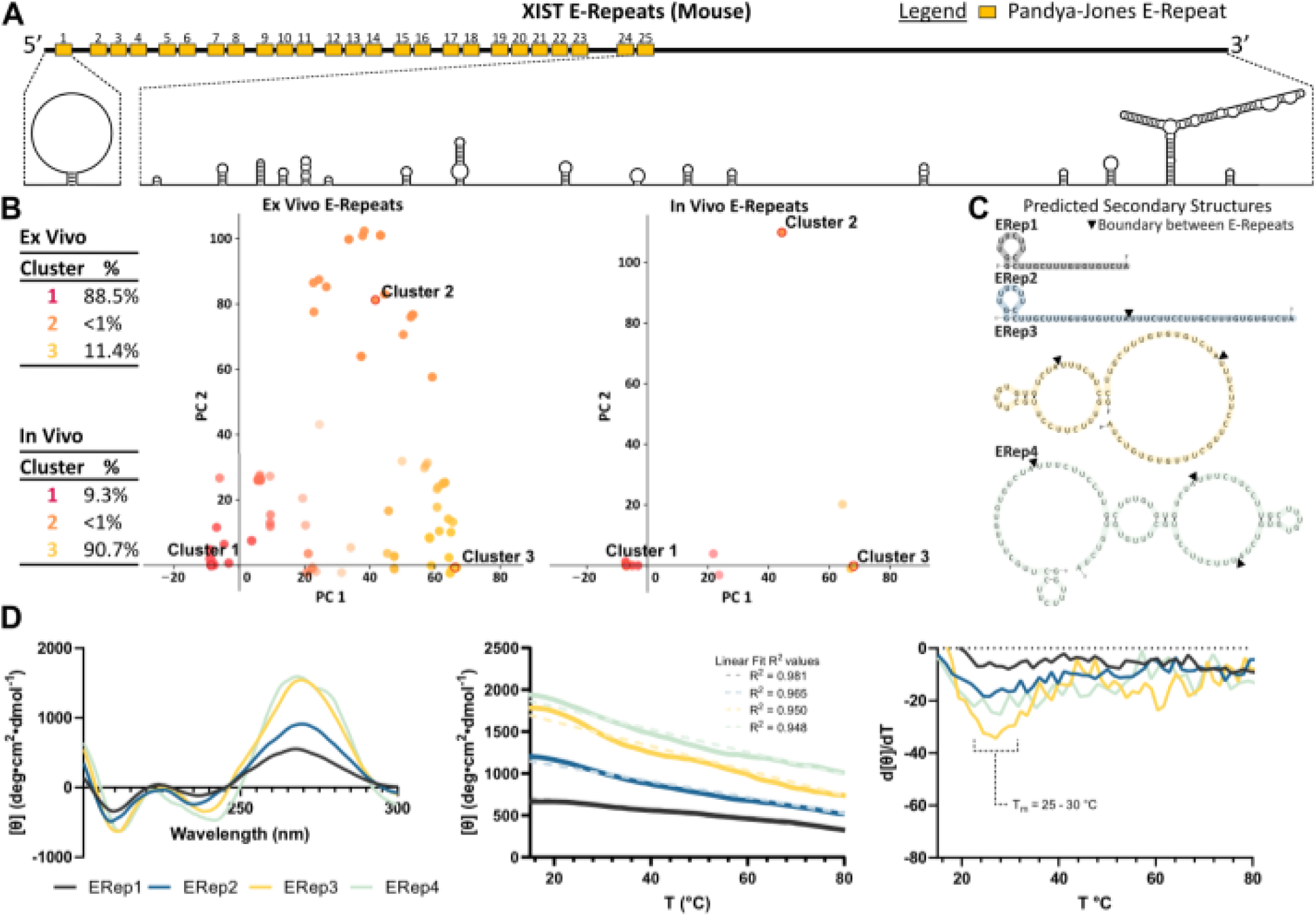
Structural analysis of the XIST E-Repeats (mouse). **A)** Linear representation of the E-Repeat domain of the XIST IncRNA and 2D structure of the E-Repeat domain. **B)** DaVinci structural analysis of the E-Repeats *ex vivo* (SRR3467748) and *in vivo* (SRR3474711). Three clusters of structures are observed under each condition; the tables on the left indicate the populations of each cluster. **C)** In silico 2D structural models of the synthetic E-Repeat constructs used in this study. **D)** CD spectroscopy spectra of the synthetic E-Repeat constructs measured at 298K (left), the melting curves 15°C to 80°C of the synthetic ERep RNAs (middle), and the first derivative of the melting curves (right). Melting temperatures are indicated in the figure.

### ELAVL1 RRM12 facilitates binding to the E-Repeats

We investigated the *in vitro* affinity of full-length ELAVL1 (ELAVL1-FL) with each synthetic E-Repeat construct (ERep1-4) using filter dot blot assays (**Figure 3A, B, Supplemental Figure 1A-D**). Quantifying the fraction of bound RNA yielded sigmoidal binding curves characteristic of cooperative interaction^45^. Fitting the data to the Hill equation revealed an apparent dissociation constant (K_D_) of 35±1 nM for a single E-Repeat, with a Hill coefficient (n_H_) of 1.5. Binding affinity remained consistently in the low nanomolar range across increasing repeat lengths (K_D_ = 61±6 for ERep2, 28±1 for ERep3, and 38±2 nM for ERep4). However, Hill coefficients increased with RNA length, shift from ∼1.5 for ERep1 and ERep2 to ∼2.3 for ERep3 and ERep4. These results indicate that while optimal affinity requires only a single E-Repeat, longer RNA templates promote higher-order assembly cooperativity.

To evaluate the ionic sensitivity of the ELAVL1-E-Repeat interface, we conducted binding assays with ERep4 across NaCl concentrations ranging from 50 mM to 300 mM (**Figure 3C**). Increasing ionic strength resulted in a moderate 2.3-fold decrease in binding affinity, shifting the K_D_ from 62±5 nM at 50 mM NaCl to 86±8 nM at 150 mM NaCl, and 146±7 nM at 300 mM NaCl. This modest reduction under high-salt conditions demonstrates that the interaction is not purely driven by electrostatic attraction, but relies significantly on non-ionic forces such as hydrogen-bonding and hydrophobic interactions.

To dissect the specific protein domains driving RNA recognition, we expressed and purified individual RRMs (RRM1, RRM2, RRM3) and tandem RRM combinations (RRM12, RRM23, and RRM123), and measured their binding affinities by filter dot blot (**Figure 3D**). Removal of the N- and C-terminal disordered regions (RRM123 construct) yielded affinities comparable to or slightly better than the full-length protein, binding ERep1, ERep2, ERep3, and ERep4 with K_D_ values of 19±2, 39±2 nM, 16±1, and 21±1 nM, respectively (**Figure 3D**). Hill coefficients for RRM123 similarly increased with repeat length (n_H_ ∼1.2 for ERep1, ∼1.5 for ERep2, and ∼1.9 for ERep3 and ERep4).

Isolated RRMs displayed significantly weaker micromolar-to-high-nanomolar binding affinities (**Figure 3D**). RRM1 bound ERep1 and ERep2 with K_D_ values of 1.29±0.22 µM and 1.08 ±0.15 µM, with affinity increasing on longer templates (K_D_ = 561±68 for ERep3 and 423±50 nM for ERep4). RRM2 exhibited the weakest binding, with no measurable binding below 1 µM (K_D_ ranges from 1.87±0.07 µM to 2.74±6.0 µM). RRM3 displayed intermediate affinities, binding ERep1 and ERep2 at 928±125 nM and 1.23±0.17 µM and ERep3 and ERep4 at 679±72 nM and 619±52 nM. No length-dependent trends in Hill coefficients were observed for individual RRMs (**Supplemental Figure 1B**).

In contrast to individual domains, the tandem RRM12 construct exhibited dramatically enhanced binding affinity and clear length dependence, binding ERep1 at 91±16 nM, ERep2 at 112±15 nM, ERep3 at 39±3 nM, and ERep4 at 44±3 nM (**Figure 3D**). Hill coefficients for RRM12 also increased with repeat length, rising from ∼1.5 (ERep1) to ∼2.2 (ERep3 and 4). The tandem RRM23 construct bound with mid-nanomolar affinity but displayed an inverse length dependence, with K_D_ values weakening from 320±21 nM (ERep1) to 545±46 nM (**Figure 3D**). Together, these domain-mapping experiments demonstrate that while all three RRMs contribute to RNA engagement, the tandem RRM12 domain serves as the primary functional unit driving high-affinity binding to E-Repeat RNA.

**Figure 3.**
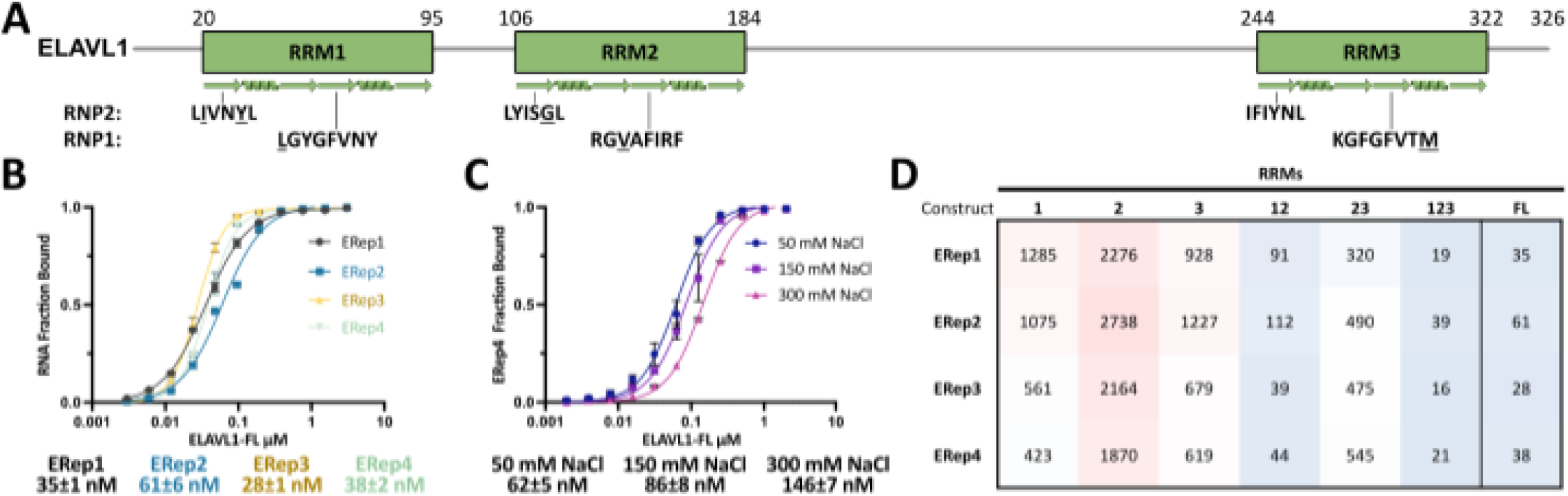
ELAVL1 binds ERep RNA with nanomolar binding affinity. **A)** Sausage plot representing the 2D structure of ELAVL1. Deviations from consensus RNP motifs are underlined. **B)** Representative filter dot blot for full length ELAVL1 with EReps 1-4; fitted plots are shown below. Protein concentration is in log 10. **C)** Salt titration binding curves gauging the ionic dependency between full length ELAVL1 and ERep4. **D)** Heat plot matrix of binding affinities between truncated ELAVL1 constructs and ERep1-4. Experiments were run in duplicate. Warm colors (red) indicate a weak interaction, whereas cooler colors (blue) indicate stronger binding affinities. See also Supplemental Figure 1 for Hill coefficients and error bars.

### PTBP1 binds the Xist E-Repeats through interactions with tandem RRMs 12 and 34

We evaluated the binding mechanics of full-length PTBP1 (PTBP1-FL) across the minimal E-Repeat constructs using filter dot blot assays (**Figure 4A, B, Supplemental Figure 2A-D**). PTBP1-FL bound a single E-Repeat with a K_D_ of 73±7 nM. Binding affinity improved progressively as the number of repeats increased, reaching 27±4 nM for ERep2, 18±2 nM for ERep3, and 14±1 nM for ERep4, demonstrating a >5-fold enhancement on the four-repeat template (**Figure 4B**). The Hill coefficient shifted from 2.3 on ERep1 to 1.5 on ERep2, remaining consistent at ∼1.6 for ERep3 and ERep4. This reduction in Hill coefficient alongside enhanced affinity suggests a shift from cooperative protein–protein assembly on short constructs to avidity-driven binding on multi-repeat templates.

To determine the electrostatic contribution to the PTBP1-E-Repeat interface, we measured binding affinities across a salt gradient from 50 mM to 300 mM NaCl with ERep4 (**Figure 4C**). Increasing ionic strength produced only a modest 2-fold decrease in binding affinity, shifting the K_D_ from 15±1 nM at 50 mM NaCl to 17±2 nM at 150 mM NaCl, and 31±3 nM at 300 mM NaCl. This high-salt stability confirms that the PTBP1-E-Repeat interaction is predominantly mediated by non-electrostatic contacts.

To map the functional domains responsible for E-Repeat recognition, we tested individual RRMs (RRM1, RRM2, RRM3, and RRM4), tandem RRMs (RRM12 and RRM34), and tri-RRM constructs (RRM123 and RRM234) (**Figure 4D**). Isolated RRM1 bound weakly to ERep1 (1.0±0.1 µM), with affinity moderately improving on ERep4 (739±75 nM). RRM2 and RRM3 displayed constant, length-independent mid-nanomolar affinities across ERep1-4 194-216 nM for RRM2; 222-271 nM for RRM3). Isolated RRM4 showed no detectable binding under the tested conditions.

Tandem RRM constructs revealed distinct binding profiles. Linking RRM3 and RRM4 (RRM34) produced a substantial gain in binding affinity that amplified with RNA length, shifting from 137±8 nM nM on ERep1 to 50±7 nM on ERep4. Conversely, the tandem RRM12 construct maintained relatively constant binding across repeat lengths (123±8 nM on ERep1 to 177±3 nM on ERep4). Extending the domain boundaries to tri-RRM constructs further improved affinity: RRM123 bound ERep1-4 with K_D_ values of 27-48 nM, while RRM234 bound with K_D_ values of 15-32 nM. Because neither tri-RRM construct fully recapitulated the 14 nM affinity to full-length PTBP1 on ERep4, these results demonstrate that maximum binding efficiency requires coupling across the complete four-RRM architecture.

**Figure 4.**
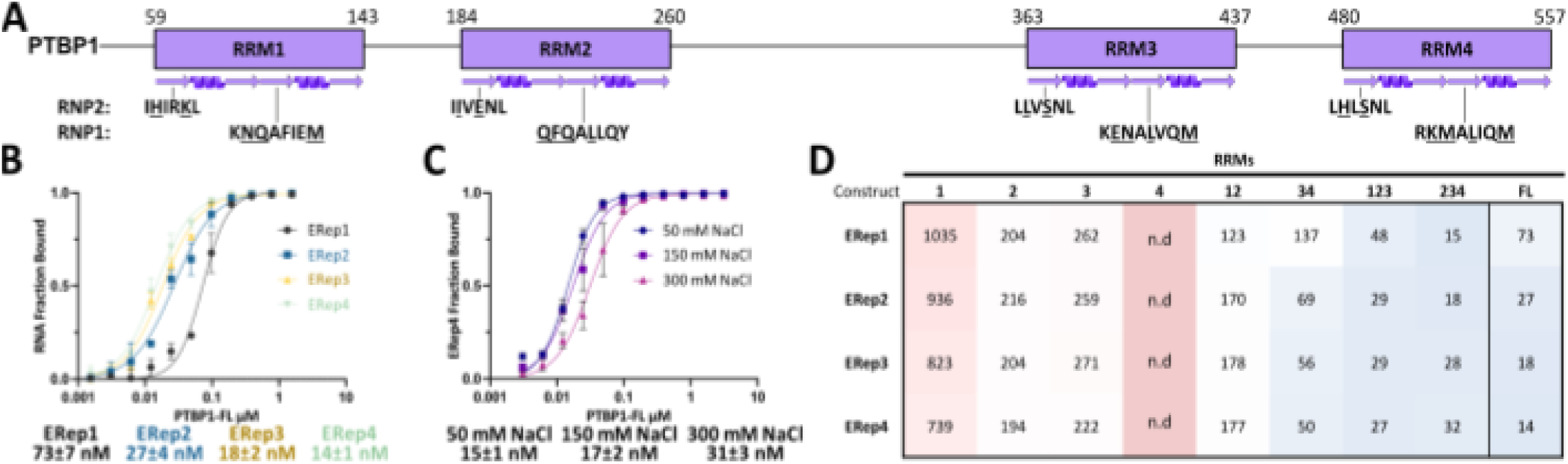
PTBP1 binds ERep RNA in a length-dependent fashion. **A)** Sausage plot of PTBP1 showcasing the secondary structures of RRMs and their corresponding RNP motifs. **B)** Representative binding curves generated from filter dot blots for full length PTBP1 with ERep1-4. **C)** Salt titration with ERep4. **D)** Heat plot matrix of binding affinities between truncated PTBP1 constructs and ERep1-4. See also Supplemental Figure 2 for Hill coefficients and error bars.

### RRM1 drives TDP43 binding to the E-Repeats

We evaluated the binding affinity of TDP43 across the synthetic E-Repeat series using filter dot blot assays, focusing on the tandem RNA-binding domain construct (TDP43 RRM12) and isolated RRMs (**Figure 5A, B, Supplemental Figure 3A-D**). TDP43 RRM12 bound the E-Repeats in a clear length-dependent manner. The dissociation constant decreased progressively from 210±30 nM on ERep1 to 134±17 nM on ERep2, 88±8 nM on ERep3, and 71±8 nM on ERep4. Over the same repeat gradient, Hill coefficients increased from 1.3 (ERep1) to 2.3 (ERep4), reflecting increasing binding cooperativity on multi-repeat constructs.

To test the ionic sensitivity of the TDP43 RRM12-E-Repeat complex, we performed salt titrations with ERep4 across a range of 50 mM to 300 mM NaCl (**Figure 5C**). TDP43 RRM12 exhibited a pronounced salt dependence: in 50 mM NaCl, RRM12 bound ERep4 with a dissociation constant of 64±2 nM; this value weakened to 249±15 nM in 150 mM NaCl, and to 754±131 nM in 300 mM NaCl. Accompanying this ∼12-fold loss in affinity, Hill coefficients decreased from 1.8 at 50 mM NaCl to 1.3 at 300 mM NaCl. This strong salt dependence indicates that electrostatic interactions contribute significantly to TDP43 RRM12 binding.

The individual RRMs, RRM1 and RRM2, have nanomolar binding affinities with the E-Repeats (**Figure 5D**). RRM1 binds ERep1 with a 203±17 nM dissociation constant and 209±13 nM with ERep2. Binding affinity increases slightly with E-Repeat length as RRM1 binds ERep3 and ERep4 at 160±12 nM and 156±8 nM, respectively. RRM2 exhibits weaker binding to the E-Repeats, only reaching a maximum of ∼25% RNA fraction bound across filter dot blots. RRM2 binds ERep1 with a K_D_ of 418±192 nM; affinity increases slightly with E-Repeat length, where RRM2 binds ERep2 with a K_D_ of 363±157 nM and ERep3 with a K_D_ of 280±41 nM. No further increase in binding affinity is seen with ERep4, which binds with a K_D_ of 313±53 nM. Because isolated RRM1 largely recapitulates the affinity of the tandem RRM12 construct, whereas RRM2 binds weakly and incompletely, these results demonstrate that RRM1 is the primary domain driving TDP43 binding to E-Repeat RNA.

**Figure 5.**
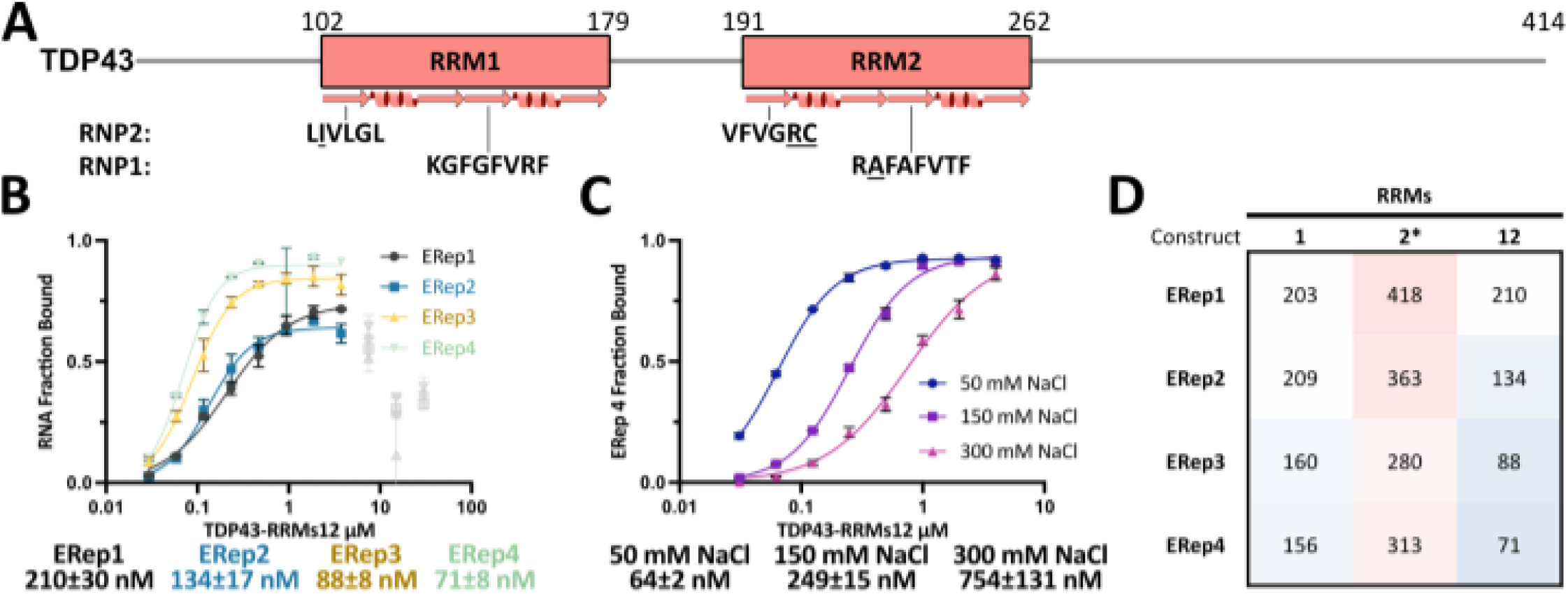
TDP43 binds ERep RNA in a length dependent fashion. **A)** TDP43 is composed of 2 RRMs. Underlined amino acids represent deviations from the consensus RNP2 and 1 sequences. **B)** Representative binding curves generated from filter dot blots for TDP43 RRMs 12 with ERep1-4 (left) and salt titration with ERep4 (right). **C)** Heat plot matrix of binding affinities between truncated TDP43 constructs and ERep1-4. *K_D_ values for RRM2 where interpolated using a fraction bound of 0.25, the maximum for this construct. See also Supplemental Figure 3 for Hill coefficients and error bars.

### hnRNPU binds the E-Repeats through its C-terminal RGG domain

We evaluated the binding interaction between hnRNPU and the synthetic E-Repeat series using filter dot blot assays (**Figure 6A, B, Supplemental Figure 4A-D**). Full-length hnRNPU (U1) bound all E-Repeat constructs with single-digit nanomolar binding affinity (K_D_ = 3.5±0.3 for ERep1, 5.17±0.5 for ERep2, 5.73±0.3 for ERep3, and 7.04±0.3 for ERep4). The Hill coefficients ranged from 1.5 to 1.9 across repeat lengths, reflecting modest binding cooperativity.

To determine which domain is primarily responsible for binding, we expressed and purified constructs spanning different regions of hnRNPU, including sub-fragments of the C-terminal RGG domain (U4 amino acid 679 - 825; U4A, amino acids 679 - 742) and constructs lacking the RGG domain (U3, U7, and U8) (**Figure 6A**). Constructs lacking the RGG domain (U3, U7, and U8) showed no detectable binding to any E-Repeat RNA (**Supplemental Figure 4D**). In contrast, the truncated RGG construct U4A bound with low-nanomolar affinity ( K_D_ = 23.9±0.4 for ERep1, 77.9±1.1 for ERep2, 51.9±0.8 for ERep3, and 45.2±2.1 nM for ERep4) (**Figure 6C**). Extending the domain boundary to include the full C-terminal RGG region (U4), the binding affinities nearly recapitulated the dissociation constants observed for the full-length hnRNP U: low nanomolar dissociation constants, between 3 and 14 nM, were observed. These results demonstrate that the extended RGG domain is the primary functional region mediating hnRNPU binding to E-Repeat RNA.

To assess the electrostatic contribution to hnRNPU binding, we performed salt titrations with full-length hnRNPU and ERep4 across NaCl concentrations from 50 mM to 300 mM (**Figure 6D**). The binding affinity between the full-length E-Repeats and hnRNPU decreased only moderately (15 to 30 nM) as the salt concentration increased (50 mM to 300 mM NaCl). This is also the case when NaCl titrations are carried out with the RGG domain U4 construct (**Supplemental Figure 4D**). Hill coefficients decreased across the salt gradient, dropping from ∼1.9 at 50 mM NaCl to ∼1.2 at 300 mM NaCl. These results demonstrate that while electrostatic interactions contribute to binding cooperativity, non-electrostatic contacts predominantly stabilize the low-nanomolar hnRNPU-E-Repeat complex.

**Figure 6.**
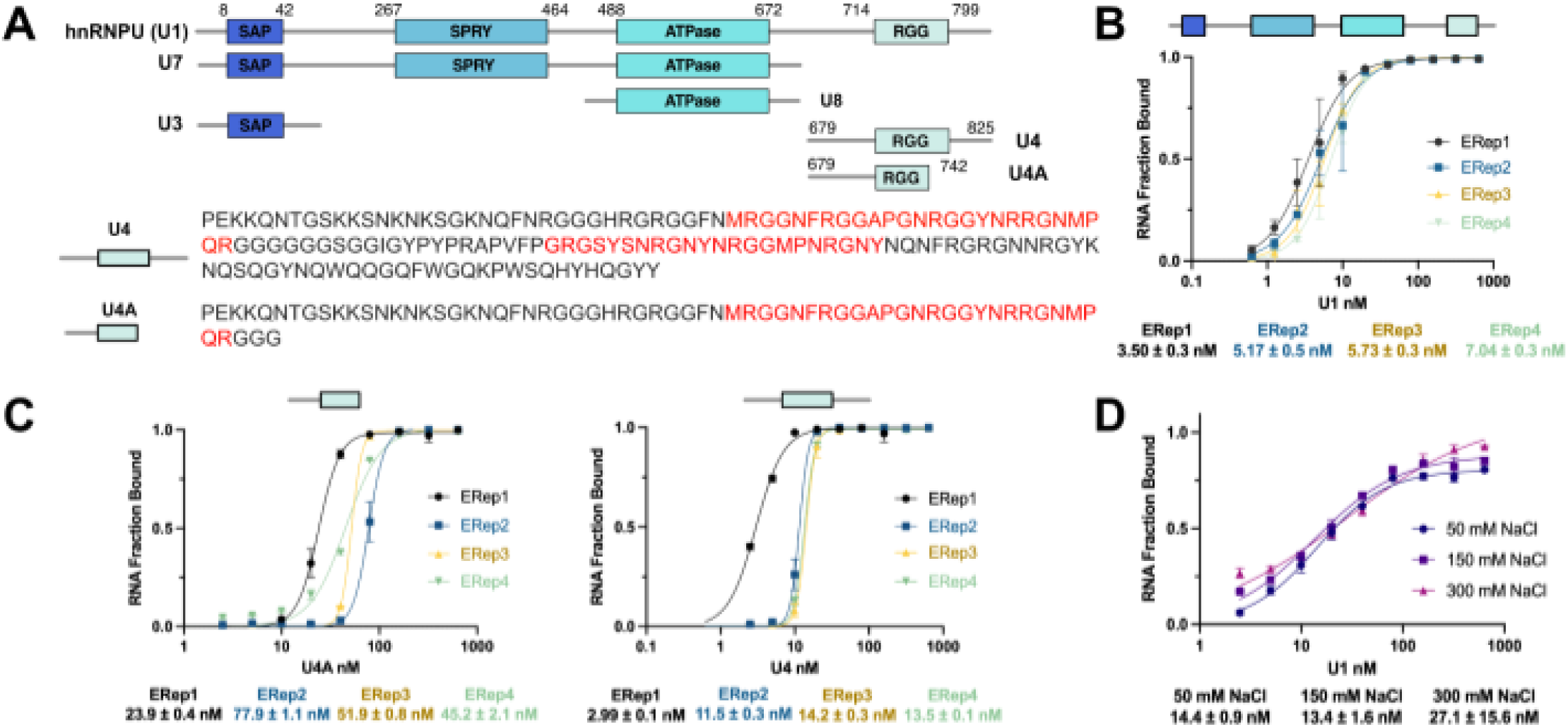
hnRNPU binds E-Repeat RNA. **A**) Schematic representation of constructs used in this study. Highlighted in red are both RGG regions in the RNA binding domain of constructs U4 and U4A. **B**) Fitted binding curves of full-length hnRNPU (U1) construct in the presence of E-Repeat RNA length 1 through 4. **C**) Fitted binding curves of hnRNPU RGG constructs U4 and U4A in the presence of E-Repeat 1 through 4. **D**) Salt gradient of U1 construct in the presence of ERep4. Values underneath graphs represent the equilibrium dissociation constant (K_D_) between the protein and the RNA indicated in the graphs. Error bars represent the standard error of the mean (SEM). Experiments were conducted in duplicates. See also Supplemental Figure 4 for Hill coefficient and error bars.

### *In vitro* analysis of higher-order protein occupancy on an E-Repeat array

Having established individual RBP binding affinities, we evaluated the propensity of E-Repeat RNA to form higher-order assemblies on ERep4 using electrophoretic mobility shift assays (EMSAs) across native agarose and PAGE gel matrices (**Figure 7A**). Single-protein titrations revealed distinct complex formation behaviours: PTBP1-FL and ELAVL1-FL formed up to four discrete, slower-migrating higher-order bands on ERep4, with gel-shift K_D_ values of 5.92±0.83 nM and 8.95±0.53 nM, respectively (**Figure 7A, Supplemental Figure 6A, B**). In contrast, TDP43 RRM12 produced a diffuse smear rather than discrete shifted bands (K_D_ = 355 nM ± 260) (**Supplemental Figure 6C**). Full-length hnRNP U formed a well-retained complex on PAGE gels (K_D_ = 9.04±1.1 nM), but did not show binding on agarose gels (**Supplemental Figure 6D**).

Using matrix conditions optimized for each protein, we performed competition EMSA titrations to monitor co-occupancy and competitive displacement between PTBP1 and the three remaining RBPs on ERep4 (**Figure 7B**). When PTBP1 is titrated into 1 µM TDP43 RRM12, PTBP1-specific complex bands emerged at concentrations as low as 3.9 nM PTBP1 (**Figure 7B**). Reciprocally, titrating TDP43 RRM12 into 250 nM PTBP1 caused a distinct lower molecular weight band to surface, gradually replacing the upper PTBP1 complex band by 1 µM TDP43 (**Figure 7B**). These shifts demonstrate that PTBP1 and TDP43 RRM12 can co-occupy ERep4 simultaneously, with TDP43 partially altering PTBP1 assembly at superstoichiometric ratios.

Co-incubation of PTBP1 and ELAVL1 revealed competitive and co-occupancy dynamics governed by molar ratios (**Figure 7B**). When PTBP1 is present in excess (1 µM) over 250 nM ELAVL1, an intermediate complex band migrates between the single-protein controls, indicating simultaneous co-occupancy on ERep4. However, when ELAVL1 was titrated into 250 nM PTBP1, ELAVL1 displaced PTBP1 starting at 31.3 nM, completely collapsing the PTBP1 double-band into an ELAVL1-like single band at 1 µM ELAVL1 (**Figure 7B**).

In contrast, co-incubation of PTBP1 and hnRNPU showed an absence of co-occupancy or competitive displacement (**Figure 7B**). Titrating PTBP1 into 250 nM hnRNPU produced PTBP1 gel shifts identical to PTBP1 alone. Reciprocally, titrating up to a 4-fold molar excess of hnRNPU (1 µM) into 250 nM PTBP1 had no effect on PTBP1 binding (**Figure 7C**). Together, these EMSA experiments demonstrate that PTBP1 forms stable, co-occupying assemblies with TDP43 RRM12 and ELAVL1, with ELAVL1 competing strongly for binding sites, whereas hnRNPU binding is completely occluded by PTBP1 on the ERep4 RNA.

**Figure 7.**
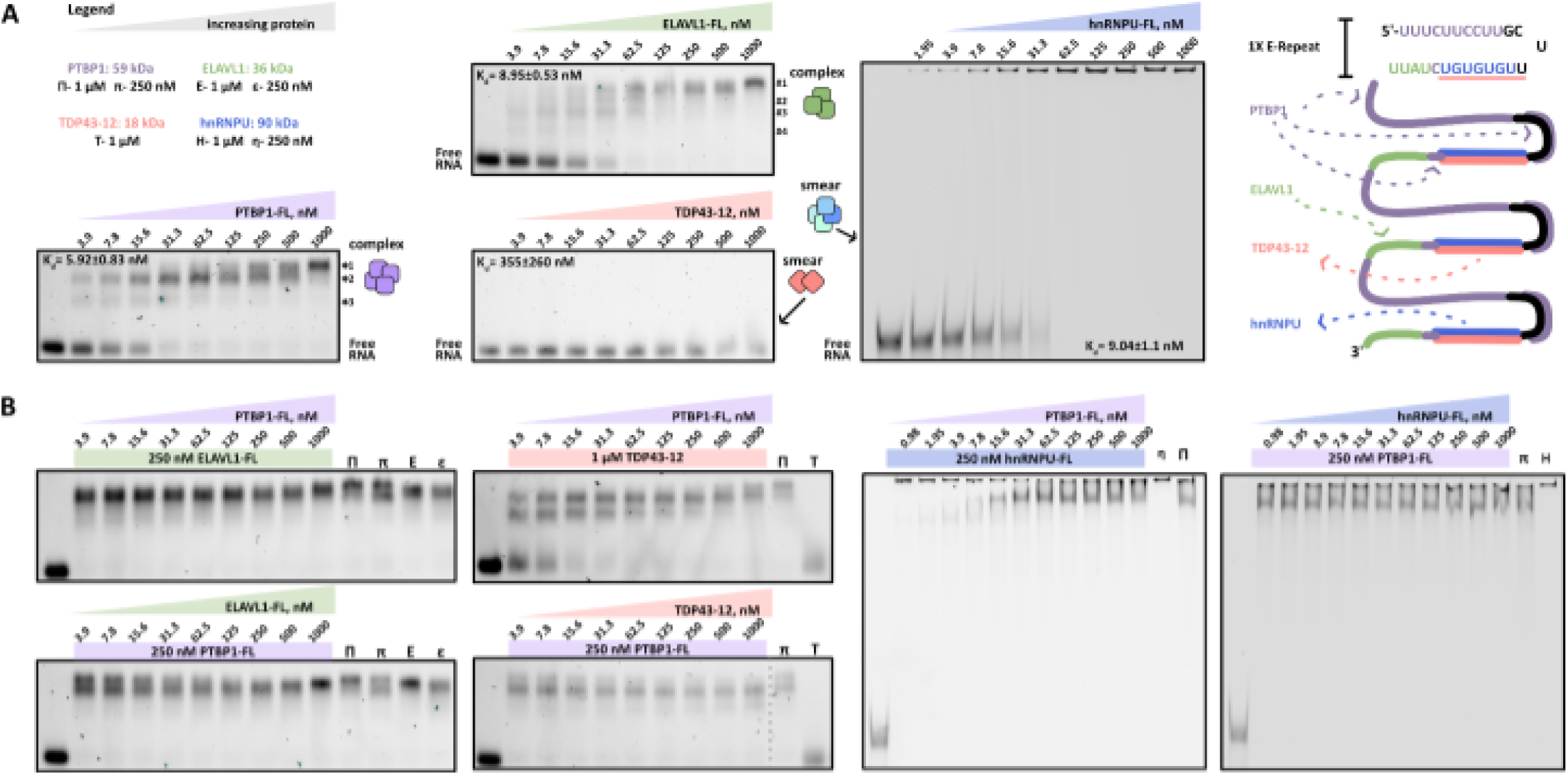
ERep4 acts as a protein landing pad with numerous binding sites. **A**) EMSAs with PTBP1, ELAVL1, TDP43 (agarose), and hnRNPU (PAGE) run against ERep4. * and # indicate separate binding events for PTBP1 and ELAVL1, respectively. The putative PTBP1 (purple), TDP43 (red), and ELAVL1 (green) binding motifs are shown on the right. **B**) Competition EMSAs between PTBP1 and ELAVL1 (left), PTBP1 and TDP43 (middle), and PTBP1 and hnRNPU (right). The vertical grey dashed line in the TDP43-PTBP1 competition assay corresponds to a crop edit (simply to fit the gels in grid format).

### *In vitro* phase separation assays reveal synergistic interactions

To examine whether E-Repeat RNA drives biomolecular condensate formation, we tested the phase separation behavior of PTBP1, ELAVL1, TDP43 RRM12, and hnRNPU in single- and multi-component reactions (**Figure 8**). In single-protein assays with ERep4 RNA under crowder-free conditions, PTBP1 underwent liquid-liquid phase separation (LLPS) in a concentration-dependent manner, forming microscopic condensates between 50 and 100 μM (**Figure 8A**). In contrast, individual incubations of ELAVL1 (5 μM), TDP43-12 (50 μM), and hnRNPU (5 μM) with the ERep4 RNA failed to form condensates under identical crowder-free conditions (**Figure 8B, Supplemental Figure 7A**). Supplemental crowding assays showed that hnRNPU formed condensates exclusively in the presence of polyethylene glycol (PEG), whereas TDP43 RRM12 did not form condensates even in the presence of PEG (**Supplemental Figure 7B**).

We next evaluated multi-protein co-incubation to determine if co-occupancy alters phase separation thresholds (**Figure 8C, D**). Co-incubation of 5 μM ELAVL1 with PTBP1 dramatically promoted phase separation, inducing condensate formation at 10 μM, a concentration at which neither protein undergoes phase separation individually (**Figure 8C, D**). This synergistic lowering of the condensation threshold was absent when ERep4 was replaced with homopolymeric polyU30 RNA (**Supplemental Figure 7C**), demonstrating that the sequence complexity of the E-Repeats is necessary for these two proteins to undergo complex coacervation. In contrast, co-incubating 5 μM hnRNPU or 10 μM TDP43 RRM12 with PTBP1 failed to promote condensate formation, even at PTBP1 concentrations up to 100 μM (**Figure 8B, Supplemental Figure 7D**). We tested the partition of hnRNPU and TDP43-12 into PTBP1 in the presence of crowder PEG; fluorescence intensity signal further confirmed that hnRNPU and TDP43 RRM12 exhibit low partitioning into PTBP1 condensates relative to ELAVL1 (**Supplemental Figure 7E**).

Finally, we tested whether E-Repeat length modulates condensate formation and partitioning by testing one to four repeat constructs at fixed protein concentrations (10 μM PTBP1, 5 μM ELAVL1) (**Figure 8D, E**). Co-incubation of 5 μM ELAVL1 or 10 μM PTBP1 with the four E-Repeat RNAs did not result in LLPS. In the ELAVL1-PTBP1 mixtures, droplet formation appeared across all repeat lengths, from ERep1 through ERep 4. Fluorescence intensity line profiles across individual condensates revealed consistent, equivalent partitioning ratios of PTBP1 and ELAVL1 across all four repeat constructs (**Figure 8E**). These results demonstrate that ELAVL1 and PTBP1 synergistically phase separate on minimal E-Repeat templates, establishing a stable co-condensate matrix that excludes transiently interacting RBPs.

**Figure 8.**
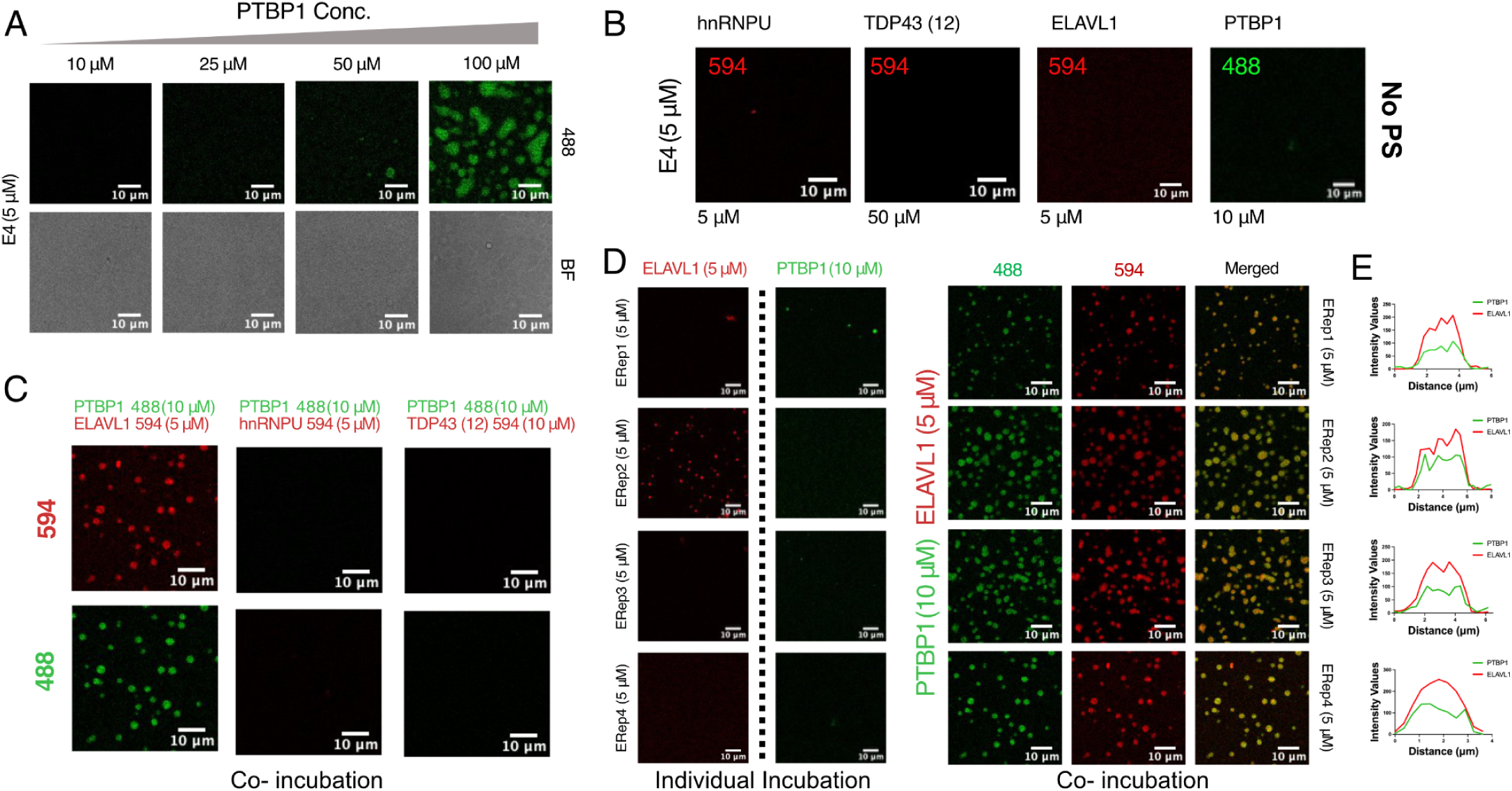
ELAVL1 and PTBP1 co-incubation enhances the phase separation propensity of both proteins. **A**) One dimensional phase diagram of PTBP1 at increasing protein concentrations in the presence of 5 μM ERep4. **B**) Individual incubation of hnRNPU, TDP43, ELAVL1, and PTBP1 in the presence of ERep4 RNA. **C**) Co-incubation of PTBP1 (Alexa 488) alternating binding partners tagged with Alexa 594 (TDP43 and hnRNPU) **D**) (left) Individual incubation of PTBP1 (Alexa 488) and ELAVL1 (Alexa 594) in the presence of increasing length of E-Rep4 RNA (right) and co-incubation of both proteins. **E**) Line profile showing intensity signal line profile of both PTBP1 (488) and ELAVLY (594).

## Discussion

LncRNAs often operate at remarkably low cellular copy numbers, raising fundamental questions about how they have outsized effects on gene expression. In this study, we provide a biophysical and biochemical dissection of the XIST E-Repeat domain and its cognate RBPs (PTBP1, ELAVL1, TDP43, and hnRNPU) to elucidate how lncRNA repetitive elements function as molecular scaffolds. We show that the E-Repeat domain exists as a dynamic, predominantly unstructured structural ensemble. Minimal single E-Repeats (26 nucleotides) are sufficient to engage all four RBPs with low nanomolar affinities, while extending repeat length up to four tandem repeats (∼98 nucleotides) increases binding cooperativity and capacity without compromising affinity. Crucially, we demonstrate that PTBP1 and ELAVL1 co-occupy multi-repeat constructs to form stable higher-order assemblies that synergistically lower the threshold for LLPS. In contrast, TDP43 and hnRNP U display more transient, displaceable binding modes that do not support co-condensation. Together, these findings demonstrate that lncRNAs can trade transcript copy number for multivalent sequence repetition, creating high-capacity binding platforms that tune protein stoichiometry and seed localized nuclear compartments^1,5,46^.

Our structural analyses and domain-mapping experiments clarify the molecular mechanisms through which these four RBPs engage the unstructured E-Repeat RNA and contextualize our quantitative affinities within established literature (**Supplemental Tables 2-5**). The deconvolution of chemical probing data revealed that while the E-Repeat domain exists as a dynamic ensemble, *in vivo* probing exhibits fewer varied structural clusters compared to *ex vivo* conditions. This structural constraint likely reflects cellular factors, including localized ionic strength, pH, and dense RBP binding, which constraint the conformational landscape of RNAs in living cells^47–50^. Observing multiple structural clusters indicates that the E-Repeat ensemble samples a diverse conformational space, a property that may be necessary to facilitate differential recognition by distinct RBPs^28,51,52^. For the multi-RRM proteins ELAVL1 and PTBP1, high-affinity binding relies on tandem domain synergy and is predominantly stabilized by non-ionic interactions. Although ELAVL1 typically recognizes AU-rich elements (AREs) in mRNA 3’ UTRs, we observed low nanomolar binding to the U-rich, low adenine E-Repeat motif^26,53^. This interaction is driven almost entirely by tandem RRM12, supporting structural models in which RRM1 acts as a primary anchor that induces a conformational reorientation, positioning the inter-domain linker and RRM2 to cooperatively clamp the RNA substrate ^54^. Similarly, PTBP1 and its tandem RRM34 construct exhibit length-dependent binding increases on multi-repeat constructs, consistent with an RNA looping mechanism where pyrimidine tracts encircle multiple RRMs to generate high avidity^55,56^. Conversely, TDP43 and hnRNPU operate through distinct structural modes. TDP43 binding is driven predominantly by RRM1; however, unlike ELAVL1 or PTBP1, its interaction displays strong salt dependence, indicating that basic surface residues framing the canonical RNP cleft provide essential electrostatic contacts that pre-anchor the RNA before base stacking occurs. Finally, hnRNPU engagement relies entirely on its intrinsically disordered C-terminal RGG domain. Truncating the second RGG cluster severely reduces binding affinity, confirming that hnRNPU binding operates via an avidity-driven network of arginine-rich repeats that accommodate single-stranded, U-rich RNA targets^31,32^.

Connecting our findings across experimental scales reveals how single-molecule binding affinities translate into higher-order assemblies and LLPS. Although equilibrium filter dot blot assays demonstrated that all four RBPs bind minimal E-Repeats with low nanomolar affinity, multi-protein EMSAs uncovered differences in complex stability and co-occupancy. PTBP1 and ELAVL1 form stable, higher order complexes on ERep4 and co-occupy the RNA when present in balanced ratios. In contrast, TDP43 and hnRNPU exhibit transient, easily displaceable interactions that fail to maintain stable co-complexes in the presence of PTPB1. This hierarchy in complex stability directly correlates with the physical nature of their binding interfaces: the salt-resilience of PTBP1 and ELAVL1 demonstrates that non-ionic hydrophobic and base-stacking interactions dominate their engagement, providing the directional crosslinking required to establish a biomolecular condensate.

Conversely, the pronounced salt sensitivity of TDP43 reflects an electrostatically dominated, highly labile interaction that cannot independently support a stable network. Consequently, this structural distinction directly governs LLPS. Consistent with prior cellular and *in vitro* reports showing that E-Repeat RNA partitions with PTBP1 into phase-separated condensates in a concentration-dependent manner to seed the inactive X-chromosome compartment, our minimal four-repeat construct faithfully recapitulates this baseline phase transition at elevated protein concentrations (50 to 100 μM). Importantly, co-incubation with ELAVL1 (5 μM) synergistically lowers the PTBP1 condensation threshold to 10 μM specifically on sequence-complex E-Repeat RNAs. Conversely, transiently interacting RBPs (TDP43 and hnRNPU) neither lower the condensation threshold nor partition efficiently into the PTBP1 matrix. This biophysical pipeline provides a compelling mechanistic rationale for cellular CLIP data, where PTBP1 and ELAVL1 densely decorate the E-Repeat region to seed active silencing foci, whereas hnRNPU and TDP43 are localized peripherally or downstream across exon 7^6,22,25,57^.

Beyond defining single-molecule and condensate mechanics, our work demonstrates that E-Repeat-mediated protein scaffolding and phase separation are intrinsic biophysical properties of the nucleoprotein complex that occur independently of DNA, chromatin components, or the broader cellular context. These findings fill a critical gap by isolating the core biochemical drivers of XIST spreading from confounding cellular variables. Nonetheless, key limitations of our *in vitro* system warrant consideration. Our minimal constructs utilize up to four tandem E-Repeats rather than the full 25-Repeat endogenous domain (∼1,500 nucleotides), and our TDP43 assays utilized a truncated RRM12 construct lacking the N- and C-terminal domains. Although these terminal domains do not significantly alter primary RNA binding affinity, they drive essential protein-protein interactions responsible for multimerization, complex stabilization, and self-directed phase separation^58^. Furthermore, while we focused on four core factors, the E-Repeat domain recruits additional intrinsically disordered RBPs, including MATR3, CELF1, and CIZ1, that contribute to cellular XIST localization and phase separation^20,21,59^. Beyond the E-Repeats, other repetitive modules across XIST, such as the B-Repeat domain and its cognate binding hnRNPK, also form condensates to facilitate cis spreading along chromatin^60^. This supports an expanded model wherein separate repetitive lncRNA domains coordinate distinct RBP networks to synergistically alter condensate morphology, optimizing transcript dispersal and gene silencing across the chromosome^5,21,60^. Future studies incorporating full-length multi-RBPs on expanded repeat arrays will further clarify how domain multivalency modulates higher-order condensate morphology. From a translational perspective, defining the minimal sequence and structural thresholds required for RBP recruitment and co-acervation establishes a framework for targeting XIST with therapeutic small molecules or oligonucleotides in X-chromosome-linked diseases. Moreover, these insights demonstrate that E-Repeat modules function as self-contained, portable scaffolding units that could be ported into synthetic lncRNA constructs to seed custom sub-nuclear compartments for cell engineering applications.

## Methods

### DaVinci analysis

Deconvolution of RNA conformations followed the DaVinci pipeline^61^. In short, the raw sequencing data generated by Smola and co-workers was downloaded from the Sequencing Read Archive. Using the SRA toolkit, paired-end reads in the form of fastq files were extracted and converted into bit vectors representing the constraint information derived from the mutational profile. The resulting constraints were fed into the CONTRAfold engine to generate a catalog of RNA structures. The collected RNA structures were then converted into dot-bracket sequences, followed by transformation into RNA secondary structure elements using rnaConvert in the Forgi package. The ensuing structure elements were subjected to dimensionality reduction via principal component analysis. *C-*means clustering was performed to cluster the PCA results, with the value of *k* being determined using the fuzzy partition coefficient.

### RNA production and purification

DNA templates, encoding for ERep1, ERep2, ERep3, and ERep4 and containing the T7 polymerase promoter sequence, were ordered from Integrated DNA Technologies. Two 5’ guanosines were added to facilitate *in vitro* transcription with the T7 polymerase. RNAs were *in vitro* transcribed using the T7 polymerase. 25 μL analytical reactions were performed to identify the optimal MgCl_2_ concentration to yield RNA, followed by large scale (mL) reactions. Briefly, analytical reactions contained 32 pmol DNA template snap cooled (95°C for 3 minutes followed by 10 minutes on ice) with equimolar T7 Top (17 nucleotide primer complementary to the T7 promoter sequence), 1.25 μL of 20X transcription buffer (100 mM Tris-HCl pH 8, 100 mM spermidine, 200 mM DTT), 2.5 μL 50% w/v polyethylene glycol (PEG) 8000, 2 μL of 100 mM each rNTP, and 1 μL of T7 polymerase (10 mg/mL). 1 M MgCl₂ was titrated to final concentrations of 20 mM, 40 mM, 60 mM, and 80 mM. Reactions were incubated at 37°C for up to 1 hour and then confirmed on a denaturing PAGE gel. *In vitro* transcription reactions were scaled up 2 mL. This mixture was then loaded onto a DNAPac™ PA200 pre-equilibrated with 6 M urea and 12.5 mM Tris-HCl pH 8.0. The RNA was eluted using an increasing gradient of 6 M Urea, 12.5 mM Tris-HCl pH 8.0, and 0.5 M NaClO4. The peaks were examined on a denaturing PAGE gel and fractions of the correct size were combined. A 1:1 ratio by volume of 100 mM NaCl was added to the eluted RNA. The suspension was then loaded onto a 100 mM NaCl equilibrated Sep-Pak column manually and dropwise. The Sep-Pak was then connected to an Akta Prime, and washed with 50 mL of water to remove urea and other salts. The RNA was then eluted with 5 mL of 50% acetonitrile. The eluent was flash-frozen and lyophilized overnight. The RNA was resuspended to the desired concentration using water.

### Cy5-Labeling

RNA was fluorescently labeled at the 3’ end by combining 100 pmol of RNA, 200 pmol pCp-Cy5 (Jena Biosciences), and 40 U T4 RNA ligase (Thermo Scientific cat. no. EL0021) in a 20 µL reaction volume containing 50 mM Tris-HCl pH 7.5, 10 mM MgCl_2_, 1 mM DTT, and 1 mM ATP, and incubated overnight at 4°C. Following this, the RNA was cleaned by passing the reaction through a G-50 column (Cytiva 27-5330-01) to eliminate excess Cy5.

### Circular Dichroism and Thermal Denaturation

All RNA was thawed on ice and kept at 4°C before measurement. 200 µL of E-Repeat RNA diluted to ∼10 µM in CD buffer (100 mM NaCl, 25 mM Na_3_PO_4_ pH 6.7) in a Hellma Analytics High Precision Cell of 1 mm path length was measured in a JASCO J-1500 CD Spectrometer. Scanning measurements from 200 to 300 nm were performed at 25°C and buffer spectra were subtracted. Spectra were then normalized to RNA concentration and smoothed by 19 points in the Savitsky-Golay smoothing algorithm in GraphPad Prism v11. For thermal denaturation experiments, the highest intensity wavelength for each E-Repeat was observed for variable temperature absorbance from 15°C to 95°C at 1°C/min with a 30s hold. Curves were normalized to RNA concentration. The first derivative of thermal denaturation data was taken with a 19 point smoothing in GraphPad Prism v11 to determine RNA melting temperatures.

### Plasmid construction

Plasmids encoding for the full-length proteins were purchased from Genscript. For protein subconstructs, protein encoding sequences were transcribed from full-length plasmids by PCR. Forward and reverse DNA primers for PCR were purchased from IDT. A double digest with KPNI and NCO1 was performed on PCR products and target pETM-11 vectors with GB1 and His-tag encoding regions. Digested DNA was then isolated by gel extraction. A quick ligation was performed using NEBlabs Quick Ligase and Quick ligase buffer. Quick ligation product was then transformed into NEB5α cells and selected by antibiotic resistance. Plasmids were extracted from NEB5α cells using the NEBiolabs miniprep protocol and sequenced by Genewiz.

### Protein expression and purification

Sequenced protein construct plasmids were transformed into BL21-DE3 competent cells. Cells were grown in LB Broth (Miller) up to OD_600_ = 0.800 before protein production was induced by addition of IPTG to a final concentration of 1 mM for 1 L LB broth. Cultures were pelleted by centrifugation (8,000 rpm, 30:00 minutes) following overnight induction. Pellets were resuspended in 25 mL resuspension/wash buffer (50 mM Tris-HCl 7.4, 300 mM NaCl, 10 mM imidazole, 5 mM βME) with supplemented DNAse I (4 µg/mL), RNAse A (4 µg/mL), lysozyme (60 µg/mL), and AEBSF-HCl protease inhibitor (1 mM) at 4°C for 10 minutes. Cells were lysed by sonication (amplitude 35, 15s on, 20s off, 3:00 duration) on ice. Cell lysate was then pelleted by centrifugation (20,000 rpm, 60:00 minutes). Flowthrough from cell lysate was passed twice through a Ni-affinity column (NEBExpress Ni Resin S1428S). The column was passed three times with wash buffer followed by two 10 mL passes with elution buffer (50 mM Tris-HCl 7.4, 300 mM NaCl, 250 mM imidazole, 5 mM βME). Protein-containing fractions were then placed in appropriately sized dialysis tubing with Tobacco Etch Virus (TEV) protease in 1 L of wash buffer (no imidazole) overnight at 4°C with mild stirring for TEV cleavage. TEV fractions were then passed twice through a Ni-affinity column, followed by 2 washes and 2 elutions. Protein-containing fractions were concentrated to 5 mL for size exclusion chromatography (SEC) on a Cytiva AKTA pure. SEC was run on a HiLoad 16/60 Superdex 75 prep grade column in the storage buffer (100 mM NaCl, 25 mM NaPO_4_., 5 mM βME). Protein containing fractions were checked by SDS-PAGE gel, concentrated and aliquoted. The hnRNPU constructs retained their His-tag and TEV cleavage site and was then purified using SEC as previously mentioned.

**Table 1.** Constructs and amino acid boundaries used in this study.

| ELAVL1 (Q15717) |  | PTBP1 (P26599) |  | TDP43 (Q13148) |  | hnRNPU |  |
| --- | --- | --- | --- | --- | --- | --- | --- |
| RRM1 | AMG+19-9<br>5 | RRM1 | AMG+59-1<br>43 | RRM1 | AMG+102-<br>200 | U8 | (His)<sub>6</sub>+TEV<br>site 1-42 |
| RRM2 | AM+106-1<br>84 | RRM2 | AMG+184-<br>260 | RRM2 | AMG+191-<br>262 | U7 | (His)<sub>6</sub>+TEV<br>site1-572 |
| RRM3 | AM+243-3<br>22 | RRM3 | AMG+363-<br>437 | RRM12 | AMG+102-<br>262 | U4 | (His)<sub>6</sub>+TEV<br>site<br>679-825 |
| RRM12 | AMG+19-1<br>84 | RRM4 | AMG+480-<br>557 |  |  | U4A | (His)<sub>6</sub>+TEV<br>site<br>679-742 |
| RRM23 | AM+106-3<br>22+GB1 | RRM12 | AMG+59-2<br>60 |  |  | U3 | (His)<sub>6</sub>+TEV<br>site<br>488-672 |
| RRM123 | AMG+19-3<br>22 | RRM34 | AMG+363-<br>557 |  |  | U1 | (His)<sub>6</sub>+TEV<br>site 1-825 |
| FL | MG+1-326 | RRM123 | AMG+59-4<br>36 |  |  |  |  |
|  |  | RRM234 | AMG+184-<br>557 |  |  |  |  |
|  |  | FL | AMG+1-55<br>7 |  |  |  |  |

### Filter dot blot (FDB) assays

Both nylon (0.45 µM; Cytiva) and nitrocellulose (0.45 µM; BioRad) membranes were soaked in a buffer solution containing 25 mM NaPO_4_. pH 7.5, 50 mM NaCl, and 5 mM BME for 30 minutes. Lower salt concentrations are used in FDB to maintain the ionic strength of the nylon membrane. Cy5-labeled RNAs were diluted to 1 nM, and then mixed with serial diluted concentrations of protein in the same buffer to a final volume of 20 µL and incubated at room temperature for five minutes. The reaction volume was then brought up to 80 µL using the respective buffer.

The filter dot blot apparatus was assembled following the instructions from the BioRad Microfiltration Apparatus Instruction Manual (cat. no. 170-6545). The membranes were first washed with 60 µL per well using the respective buffer and setting the flow valve to a gentle vacuum setting. While loading 70 µL of the samples, the flow valve was set to be exposed to air. Following, the flow valve was set to be exposed to vacuum. Once all samples were drained from the wells, the wells and membranes were washed with 70 µL of the respective buffer while still exposed to vacuum. Afterwards, the apparatus was disassembled, with the flow valve set to air exposure, and the membranes were left to dry on aluminum foil. The membranes were then imaged on BioRad ChemiDoc Touch Imaging System on a Blot/UV/Stain-Free Sample Tray, using the Cy5 Fluorophore (emission wavelength = 670 nm). Dot intensity from both membranes were quantified using ImageJ, and binding affinities were calculated in Prism using the specific hill graph fit 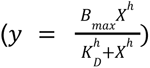.

### Electrophoretic mobility shift assays (EMSA)

EMSAs were performed in either 1% agarose or 6.5% PAGE. Agarose gel solutions were prepared by mixing 0.09 g Synergel and 0.41 g agarose in 1 mL of 100% EtOH and dissolving in 50 mL of running buffer (1xTB Buffer). 50 mL of solution was cast into 15 x 10 cm trays for optimal gel thickness. Non-competitive EMSAs were prepared via serial dilution of 10 µL concentrated protein into EMSA buffer (80 mM KCl, 20 mM HEPES pH 7.5, 6 mM MgCl_2_, 1 mM DTT, 0.02% v/v Triton, 20% glycerol) to achieve a final maximum concentration of 1 µM in 12 µL. 10 µL protein dilutions were incubated for 8 minutes at room temperature before addition of 120 pmol of Cy5-labelled E-Repeat 4 RNA. Protein-RNA binding solutions were incubated at room temperature for 15 minutes at room temperature, followed by 5 minutes on ice before addition to wells. Agarose gels were run in 1x TB at 4°C with a constant voltage of 60 V for 2 hours. PAGE gels were run in 1x TB at constant 5 W for 40-45 minutes at room temperature. Following run completion, all EMSAs were immediately covered and Cy5 imaged with a BioRad ChemiDoc Touch Imaging System on a Blot/UV/Stain-Free Sample Tray PAGE. Competition EMSAs followed the same procedure as both proteins were incubated together in the initial 8 minutes, followed by addition of the RNA. Band intensity was quantified using ImageJ, and binding affinities were calculated using Prism.

### *In vitro* phase separation assays and protein labelling

Protein constructs were labeled with either Alexa Fluor™ 488 or 594 NHS Ester (Invitrogen™, A20000) as per manufacturing instructions. Coverslips (Thorlabs) with custom-made wells of approximately 6 µL in volume were used for incubation and imaging. The condensate buffer for all proteins was composed of 100 mM NaCl, 25 mM NaPO_4_. 0.5-3% labeled protein species was used for each sample. Condensates were induced by diluting the protein of interest (PTBP1, ELAVL1, hnRNPU, and TDP43) to its desired final concentration in the assay buffer and adding RNA to induce the formation of droplets. Samples were incubated between 30 min to 1 hour at room temperature prior to imaging unless otherwise specified. Confocal images were taken on a Leica TCS SP8 X Laser Confocal Microscope equipped with a 63×/1.30 NA PLAN APO objective, using a white-light laser source and a Leica DFC310 digital color camera (2.7693 pixels/µm resolution). Images were processed and analyzed using Fiji and line profiles were plotted using Prism.

## Author Contributions

Conceptualization of paper, A.N.J., K.A.H., B.E.; Experimental design and execution of methodologies, B.E., A.K.M, A.V.C.; Data analysis and visualization, A.N.J, B.E., A.K.M., A.V.C.; Protein purification R.A.M., E.B., A.V.C., A.K.M., B.E.; Funding acquisition A.N.J.; Writing manuscript A.N.J, B.E., A.K.M., A.V.C.; Reviewing manuscript, all authors.

## Acknowledgements

The authors acknowledge the NYU Department of Chemistry Shared Instrumentation Facility, the shared facilities provided through the Materials Research Science & Engineering Center (MRSEC) program of the NSF under Award Number DMR-1420073. We thank Harsha Venkatesh for assistance with DaVinCi analysis and K means fuzzy clustering, and the members of the Jones and Haynes labs for their insights and feedback on this manuscript. A.N.J. is supported by NSF 224667 (Ideas Lab).

