## Supplemental Information for "Multivalent Scaffolding by XIST E-Repeats Tunes RNA Binding Protein Cooperativity and Lowers the Phase Separation Threshold"

### Supplemental Figures

**A**

| | RRM1 | RRM2 | RRM3 | RRM12 | RRM23 | RRM123 | FL |
|---|---|---|---|---|---|---|---|
| ERep1 | 1285±220 | 2276±2508 | 928±125 | 91±16 | 320±21 | 19±2 | 35±1 |
| ERep2 | 1075±152 | 2738±5960 | 1227±171 | 112±15 | 490±42 | 39±2 | 61±6 |
| ERep3 | 561±68 | 2164±123 | 679±72 | 39±3 | 475±46 | 16±1 | 28±1 |
| ERep4 | 423±50 | 1870±71 | 619±52 | 44±3 | 545±46 | 21±1 | 38±2 |

**B**

| | RRM1 | RRM2 | RRM3 | RRM12 | RRM23 | RRM123 | FL |
|---|---|---|---|---|---|---|---|
| ERep1 | 1.707 | n.d | 1.470 | 1.475 | 1.920 | 1.176 | 1.509 |
| ERep2 | 1.499 | 4.675 | 1.443 | 1.167 | 1.770 | 1.539 | 1.527 |
| ERep3 | 1.692 | 3.077 | 1.687 | 2.150 | 1.613 | 1.914 | 2.421 |
| ERep4 | 2.161 | 3.925 | 2.202 | 2.199 | 1.590 | 1.883 | 2.275 |

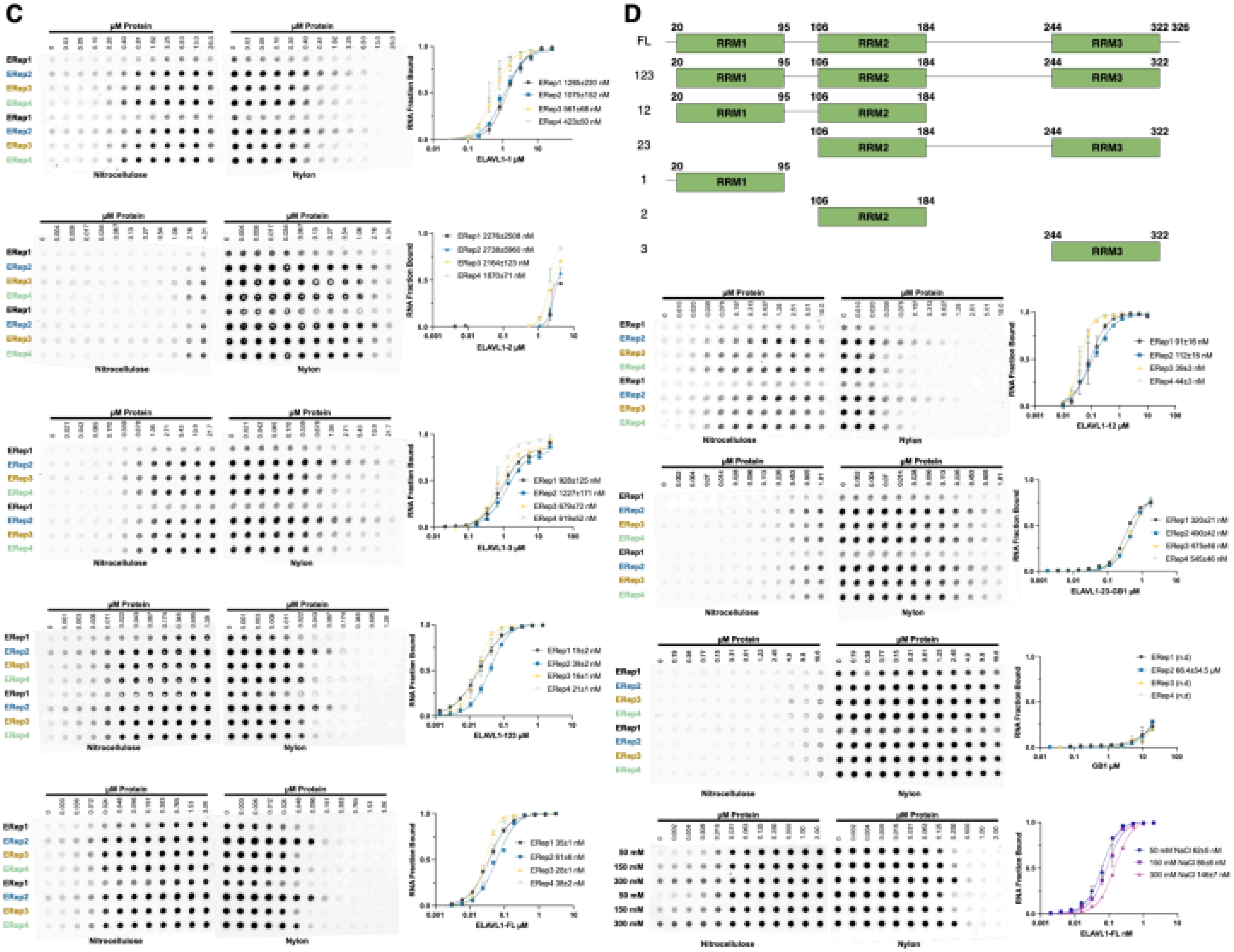


**Supplemental Figure 1**: Complete binding data for ELAVL1. **A)** Heat plot of ELAVL1 binding affinity ($K_D$) with 95% confidence interval values. **B)** Heat plot of calculated Hill coefficients for each ELAVL1 binding curve. **C)** Replicate (n=2) filter dot blots and associated binding curves for each ELAVL1 construct and salt titration. **D)** Sausage plot of ELAVL1 constructs used in this study.

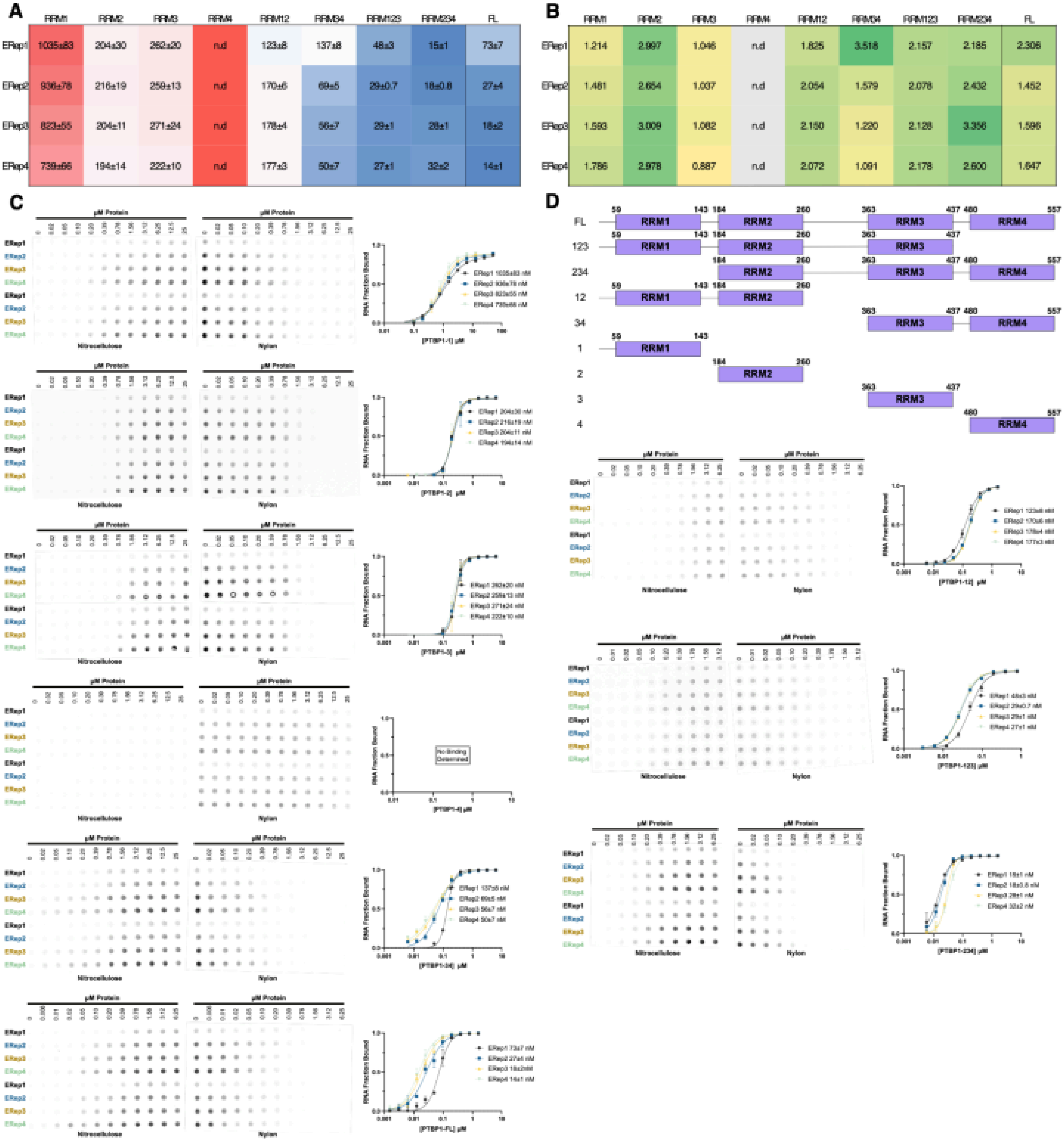


**Supplemental Figure 2:** Complete binding data for PTBP1. **A)** Heat plot of PTBP1 binding affinity ($K_D$) with 95% confidence interval values. **B)** Heat plot of calculated Hill coefficients for each PTBP1 binding curve. **C)** Replicate (n=2) filter dot blots and associated binding curves for each PTBP1 construct and salt titration. **D)** Sausage plot of PTBP1 constructs used in this study.

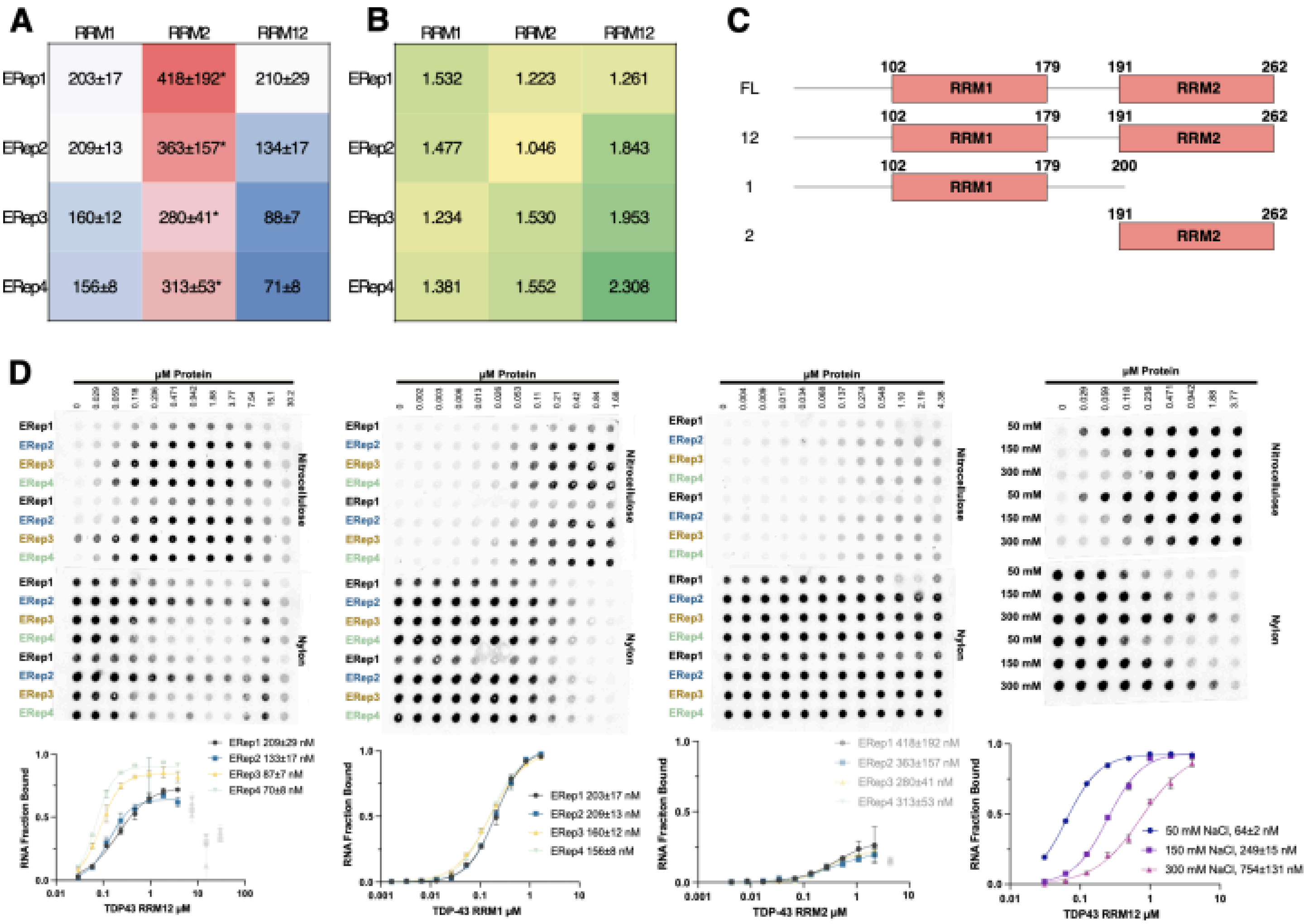


**Supplemental Figure 3:** Complete binding data for TDP43. **A)** Heat plot of TDP43 binding affinity ($K_D$) with 95% confidence interval values. **B)** Heat plot of calculated Hill coefficients for each TDP43 binding curve. **C)** Replicate (n=2) filter dot blots and associated binding curves for each TDP43 construct and salt titration. **D)** Sausage plot of TDP43 constructs used in this study.

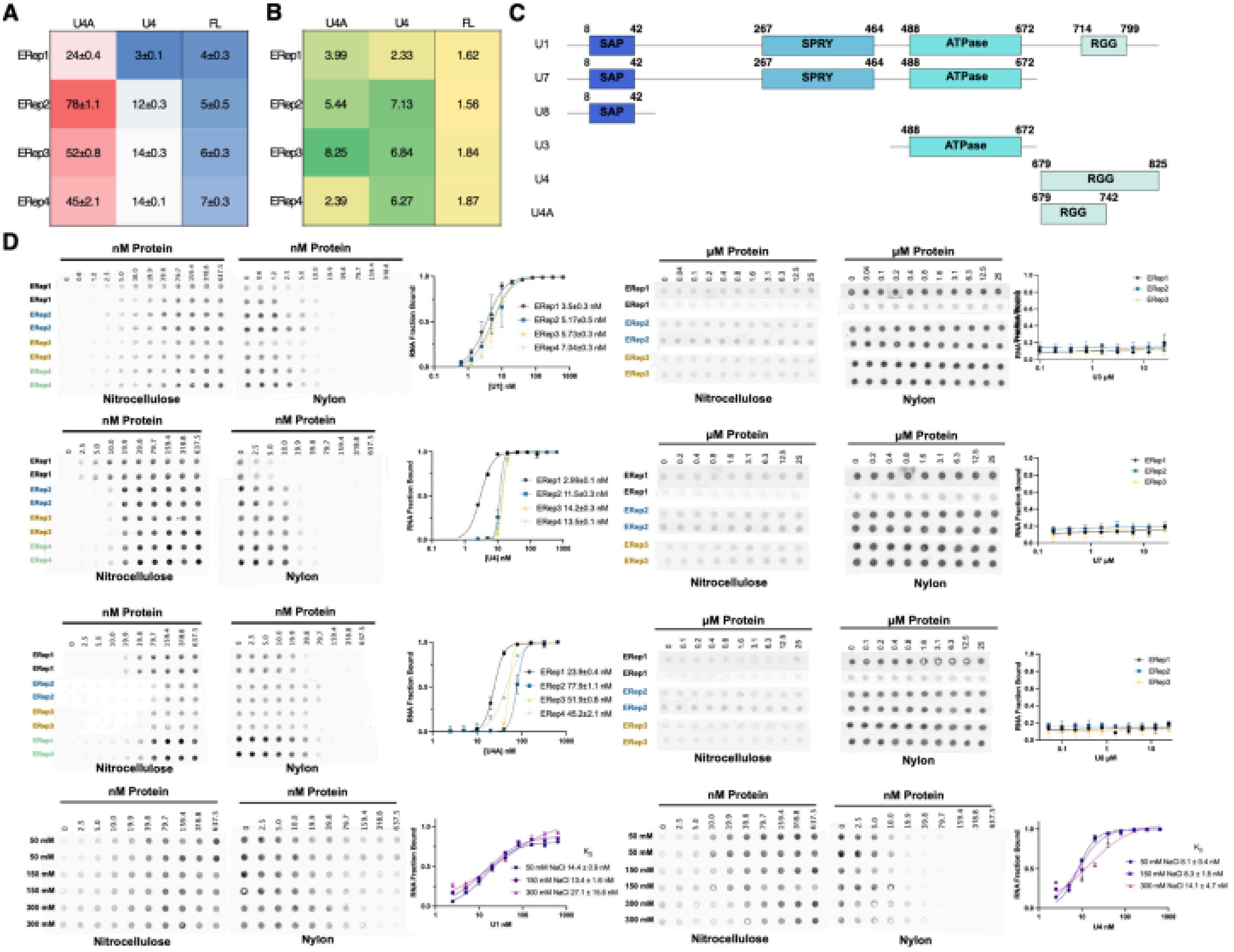


**Supplemental Figure 4:** Complete binding data for hnRNPU. **A)** Heat plot of hnRNPU binding affinity ($K_D$) with 95% confidence interval values. **B)** Heat plot of calculated Hill coefficients for each hnRNPU binding curve. **C)** Replicate (n=2) filter dot blots and associated binding curves for each hnRNPU construct and salt titration. **D)** Sausage plot of hnRNPU constructs used in this study.

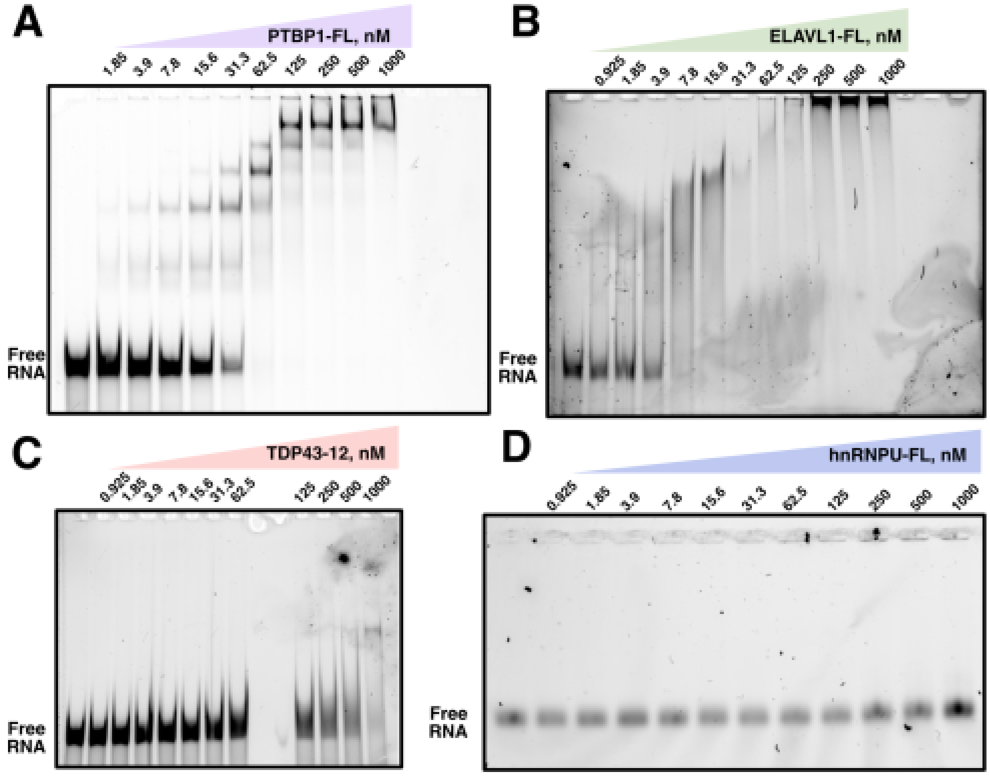


**Supplemental Figure 5:** ERep4 EMSA controls with **A)** PTBP1 (on a 6.5% PAGE gel), **B)** ELAVL1 (on a 6.5% PAGE gel), **C)** TDP43 (on a 6.5% PAGE gel), and **D)** hnRNPU (on a 1% agarose gel).

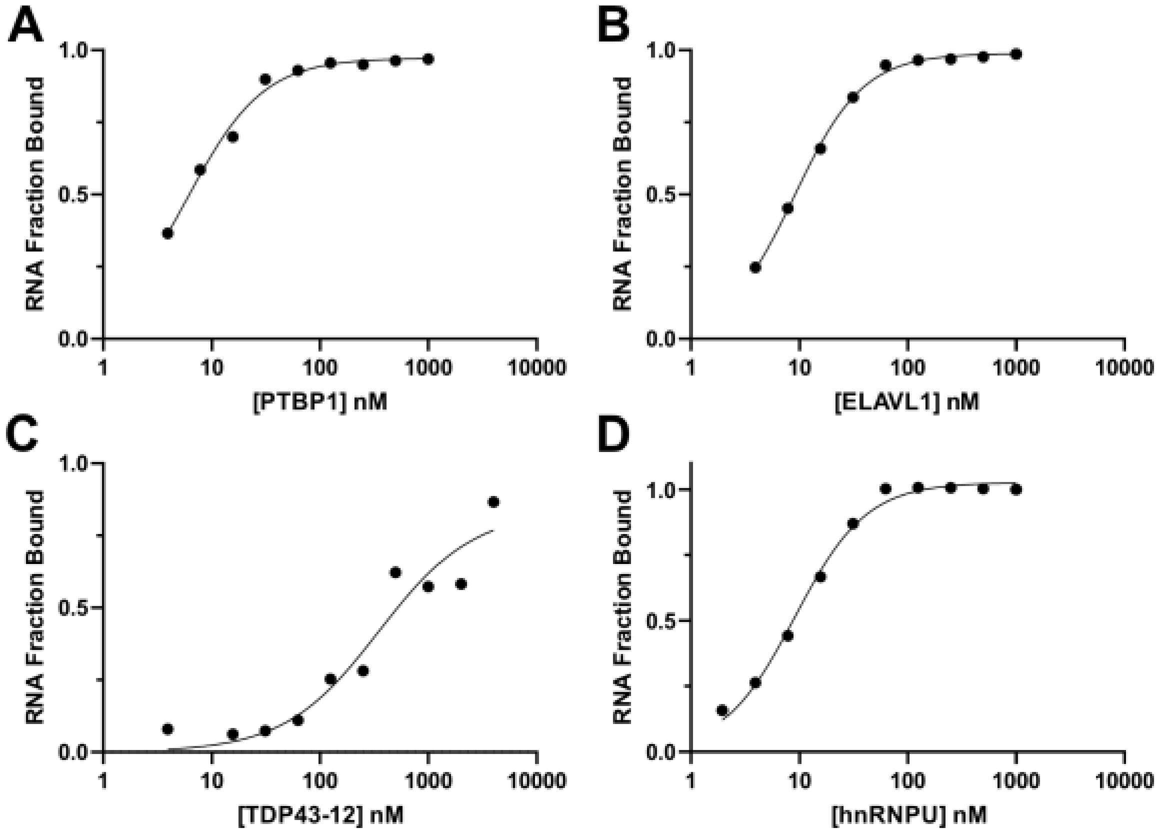


**Supplemental Figure 6:** Binding curves generated from EMSAs reported in main Figure 7B for **A**) PTBP1, **B**) ELAVL1, **C**) TDP43, and **D**) hnRNPU.

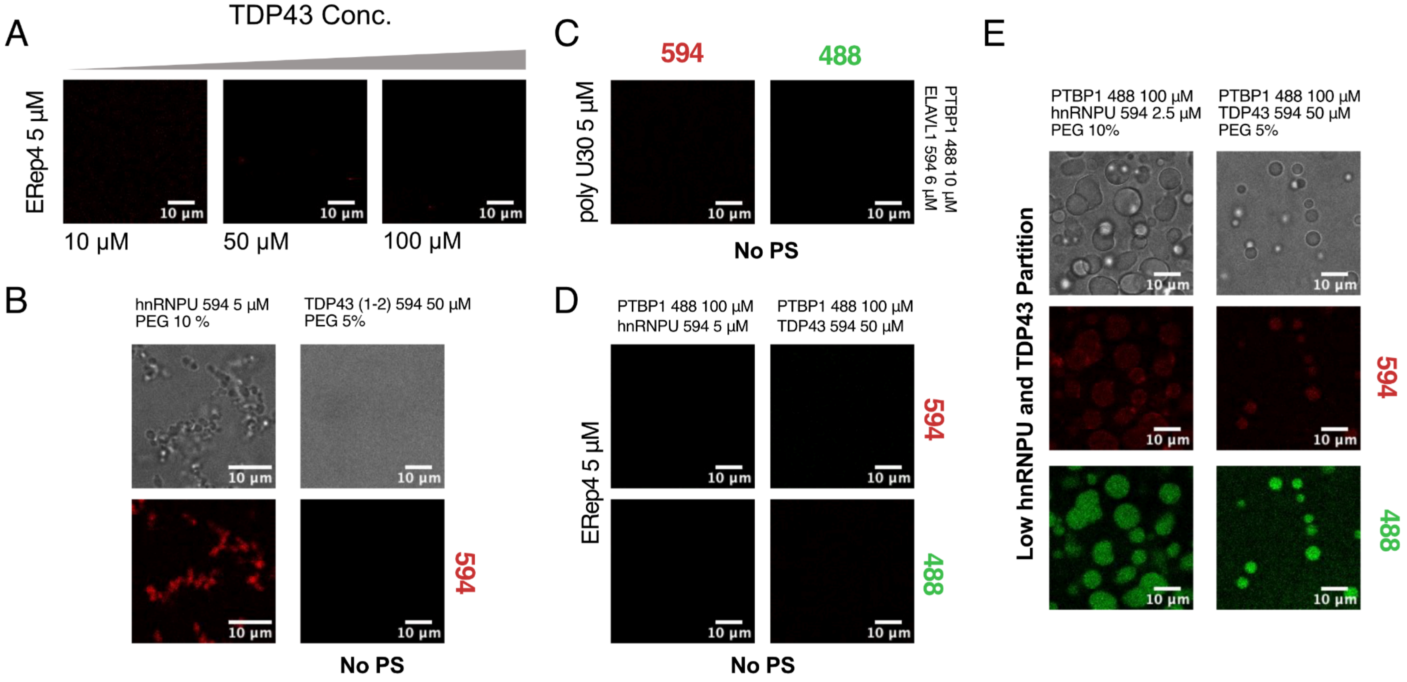


**Supplemental Figure 7:** Condensate formation controls. **A)** TDP43-12 concentration range in the presence of 5 µM ERep4 RNA **B)** hnRNPU (5 µM) and TDP43-12 (50 µM) in the presence of 5%-10% crowder PEG 8000. **C)** PTBP1 (488) and ELAVL1 (594) co-incubation in the presence of polyU30 RNA (5 µM). **D)** Co-incubation of PTBP1 (100 µM) in the presence of either hnRNPU (5 µM) and TDP43-12 (5 µM). **E)** Co-incubation of PTBP1 (100 µM) with both hnRNPU and TDP43 in the presence of 5% crowder PEG 8000.

### Supplemental Tables

**Supplemental Table 1.** Chemical probing datasets used for deconvolution analysis.

| Dataset | Chemical Probe | Coverage | Reference |
|---|---|---|---|
| Ex Vivo (SRR3467748) | 1M7 | 6.6M reads | Smola, MJ. et al. PNAS. (2016) |
| In cell (SRR3474711) | 1M7 | 1.7M reads | Smola, MJ. et al. PNAS. (2016) |

**Supplemental Table 2.** ELAVL1 Construct Binding affinities reported in the literature:

| Construct | RNA | $K_D$ | [Salt] | Incubation Time | Method | Citation |
|---|---|---|---|---|---|---|
| RRM1 | 11 base AU rich | 4.88 μM | 200 mM NaCl | Not reported | FP | (Wang et al. 2013) |
| RRM2 | 11 base AU rich | Not detectable | 200 mM NaCl | Not reported | FP | (Wang et al. 2013) |
| RRM12 | 11 base AU rich | 169 nM | 200 mM NaCl | Not reported | FP | (Wang et al. 2013) |
| RRM12 | Ar 1-51 | 81 nM | 14 mM KCl | 30 min | EMSA | (Barker et al. 2012) |
| RRM12 | cFOS-ARE | 86 nM | 14 mM KCl | 30 min | EMSA | (Barker et al. 2012) |
| RRM12 | TNFa-ARE | 150 nM | 14 mM KCl | 30 min | EMSA | (Barker et al. 2012) |
| FL | Ar 1-51 | 43 nM | 14 mM KCl | 30 min | EMSA | (Barker et al. 2012) |
| FL | cFOS-ARE | 26 nM | 14 mM KCl | 30 min | EMSA | (Barker et al. 2012) |
| FL | TNFa-ARE | 52 nM | 14 mM KCl | 30 min | EMSA | (Barker et al. 2012) |
| RRM3 | 25 mer U-rich | 61 μM | Not reported | Not Reported | CD | (Scheiba et al. 2014) |
| RRM3 | 25 mer AU-rich | 124 μM | Not reported | Not Reported | CD | (Scheiba et al. 2014) |
| FL | IL-8 (3'UTR) | 1.8 nM | 100 mM NaCl<br>20 mM KCl | 30 min | ELISA | (Nabors et al. 2001) |
| FL | VEGF (3'UTR) | 3.4 nM | 100 mM NaCl<br>20 mM KCl | 30 min | ELISA | (Nabors et al. 2001) |
| FL | COX2 (3'UTR) | 3.1 nM | 100 mM NaCl<br>20 mM KCl | 30 min | ELISA | (Nabors et al. 2001) |
| FL | TGFB (3'UTR) | 2.3 nM | 100 mM NaCl<br>20 mM KCl | 30 min | ELISA | (Nabors et al. 2001) |
| FL | TNF (3'UTR) | 18.3 nM | 100 mM NaCl<br>20 mM KCl | 30 min | ELISA | (Nabors et al. 2001) |
| FL | IL-6 (3'UTR) | 10.6 nM | 100 mM NaCl<br>20 mM KCl | 30 min | ELISA | (Nabors et al. 2001) |
| 12 | U-rich28mer | 33 nM | 150 mM NaCl | Not reported | SPR | (Kim et al. 2011) |
| 12 | A-rich 28mer | 502 nM | 150 mM NaCl | Not reported | SPR | (Kim et al. 2011) |
| 123 | U-rich28mer | 50 pM | 150 mM NaCl | Not reported | SPR | (Kim et al. 2011) |
| 123 | A-rich 28mer | 190 pM | 150 mM NaCl | Not reported | SPR | (Kim et al. 2011) |

| | | | | | | |
|---|---|---|---|---|---|---|
| 12 | poly-U 8mer | 4.7 μM | 150 mM NaCl | Not reported | SPR | (Kim et al. 2011) |
| 12 | poly-U 13mer | 195 nM | 150 mM NaCl | Not reported | SPR | (Kim et al. 2011) |
| 12 | poly-U 17mer | 40 nM | 150 mM NaCl | Not reported | SPR | (Kim et al. 2011) |
| FL | ARE 11mer(TNFa) | 82 nM | 50 mM KCl | Not reported | EMSA | (Fialcowitz-White et al. 2007) |
| FL | ARE 13mer(TNFa) | 8.9 nM | 50 mM KCl | Not reported | EMSA | (Fialcowitz-White et al. 2007) |
| FL | ARE 14mer(TNFa) | 4.8 nM | 50 mM KCl | Not reported | EMSA | (Fialcowitz-White et al. 2007) |
| FL | ARE 15mer(TNFa) | 2.5 nM | 50 mM KCl | Not reported | EMSA | (Fialcowitz-White et al. 2007) |
| FL | ARE 16mer(TNFa) | 2.5 nM | 50 mM KCl | Not reported | EMSA | (Fialcowitz-White et al. 2007) |
| FL | ARE 18mer(TNFa) | 0.83 nM | 50 mM KCl | Not reported | EMSA | (Fialcowitz-White et al. 2007) |
| FL | ARE 20mer(TNFa) | 0.44 nM | 50 mM KCl | Not reported | EMSA | (Fialcowitz-White et al. 2007) |
| FL | ARE 38mer(TNFa) | 2.0 nM | 50 mM KCl | Not reported | FA | (Fialcowitz-White et al. 2007) |
| RRM12+hinge | ARE 11mer(TNFa) | 26.2 nM | 50 mM KCl | 60 mins | FA | (Fialcowitz-White et al. 2007) |
| RRM12+hinge | ARE 13mer(TNFa) | 10.1 nM | 50 mM KCl | 60 mins | FA | (Fialcowitz-White et al. 2007) |
| RRM12+hinge | ARE 14mer(TNFa) | 5.3 nM | 50 mM KCl | 60 mins | FA | (Fialcowitz-White et al. 2007) |
| RRM12+hinge | ARE 15mer(TNFa) | 4.4 nM | 50 mM KCl | 60 mins | FA | (Fialcowitz-White et al. 2007) |
| RRM12+hinge | ARE 16mer(TNFa) | 3.5 nM | 50 mM KCl | 60 mins | FA | (Fialcowitz-White et al. 2007) |
| RRM12+hinge | ARE 18mer(TNFa) | 2.2 nM | 50 mM KCl | 60 mins | FA | (Fialcowitz-White et al. 2007) |
| RRM12+hinge | ARE 20mer(TNFa) | 1.6 nM | 50 mM KCl | 60 mins | FA | (Fialcowitz-White et al. 2007) |
| RRM12+hinge | ARE 38mer(TNFa) | 1.7 nM | 50 mM KCl | 60 mins | FA | (Fialcowitz-White et al. 2007) |
| RRM12 | ARE 13mer(TNFa) | 64 nM | 50 mM KCl | 60 mins | FA | (Fialcowitz-White et al. 2007) |
| RRM12 | ARE 16mer(TNFa) | 99 nM | 50 mM KCl | 60 mins | FA | (Fialcowitz-White et al. 2007) |
| RRM12 | ARE 20mer(TNFa) | 65 nM | 50 mM KCl | 60 mins | FA | (Fialcowitz-White et al. 2007) |
| RRM12 | ARE 38mer(TNFa) | 98 nM | 50 mM KCl | 60 mins | FA | (Fialcowitz-White et al. 2007) |

**Supplemental Table 3.** PTBP1 Construct Binding affinities reported in the literature:

| Construct | RNA | $K_D$ | [Salt] | Incubation Time | Method | Citation |
|---|---|---|---|---|---|---|
| FL | 150PYWT | 313 nM | 100 mM NaCl | Not reported | EMSA | (Taylor et al. 2022) |
| FL | CPA1 | None detected | 100 mM NaCl | Not reported | EMSA | (Taylor et al. 2022) |
| FL | CPA2 | 79 nM | 100 mM NaCl | Not reported | EMSA | (Taylor et al. 2022) |
| FL | CPA2MUT | 180 nM | 100 mM NaCl | Not reported | EMSA | (Taylor et al. 2022) |
| FL | FRAGT1 | None detected | 100 mM NaCl | Not reported | EMSA | (Taylor et al. 2022) |
| FL | FRAGT2 | None detected | 100 mM NaCl | Not reported | EMSA | (Taylor et al. 2022) |
| FL | FRAGT3 | 442 nM | 100 mM NaCl | Not reported | EMSA | (Taylor et al. 2022) |
| FL | FRAGT4 | 453 nM | 100 mM NaCl | Not reported | EMSA | (Taylor et al. 2022) |
| FL | FRAGT5 | None detected | 100 mM NaCl | Not reported | EMSA | (Taylor et al. 2022) |
| FL | FRAGT6 | 114 nM | 100 mM NaCl | Not reported | EMSA | (Taylor et al. 2022) |
| FL | FRAGT6MUT | 723 nM | 100 mM NaCl | Not reported | EMSA | (Taylor et al. 2022) |
| FL | FRAGT7 | None detected | 100 mM NaCl | Not reported | EMSA | (Taylor et al. 2022) |
| FL | PV dV-FL WT | 41 nM | 80 mM KCl | 24 mins | FA | (Avanzino et al. 2018) |
| FL | PV dV-FL WT + 4G | 63 nM | 80 mM KCl | 24 mins | FA | (Avanzino et al. 2018) |
| FL | PV dV-FL Sabin 1 | 72 nM | 80 mM KCl | 24 mins | FA | (Avanzino et al. 2018) |
| FL | PV dV-FL Sabin 2 | 69 nM | 80 mM KCl | 24 mins | FA | (Avanzino et al. 2018) |
| FL | PV dV-FL Sabin 3 | 162 nM | 80 mM KCl | 24 mins | FA | (Avanzino et al. 2018) |
| FL | PV dV-FL Sabin 3 + 4G | 167 nM | 80 mM KCl | 24 mins | FA | (Avanzino et al. 2018) |
| FL (no cys) | FL EMCV probe | 33 nM | 100 mM KCl | 20 mins | FDB | (Kafasla et al. 2011) |
| RBD 1 mutant | FL EMCV probe | 46 nM | 100 mM KCl | 20 mins | FDB | (Kafasla et al. 2011) |
| RBD 2 mutant | FL EMCV probe | 65 nM | 100 mM KCl | 20 mins | FDB | (Kafasla et al. 2011) |
| RBD 3 mutant | FL EMCV probe | 43 nM | 100 mM KCl | 20 mins | FDB | (Kafasla et al. 2011) |
| RBD 4 mutant | FL EMCV probe | 34 nM | 100 mM KCl | 20 mins | FDB | (Kafasla et al. 2011) |
| his-PTB34s | EMCV IRES Domain 1 | 83 nM | 100 mM KCl | 15 min | FDB | (Yuan et al. 2002) |
| his-PTB34s H411A/Q412A | EMCV IRES Domain 1 | 140 nM | 100 mM KCl | 15 min | FDB | (Yuan et al. 2002) |
| his-PTB34s R418A | EMCV IRES Domain 1 | 200 nM | 100 mM KCl | 15 min | FDB | (Yuan et al. 2002) |
| his-PTB34s Q424A | EMCV IRES Domain 1 | 90 nM | 100 mM KCl | 15 min | FDB | (Yuan et al. 2002) |

| | | | | | | |
|---|---|---|---|---|---|---|
| PTBP1-12 | CUUCUCUCU | 5-10 μM | Not reported | Titration | NMR | (Simpson et al. 2004) |
| FL | AGCCUCUCCUU CUCUCUGCUUC UCUCUCGCUGG CCCUU | 50 nM | none | 8 min | EMSA | (Amir-Ahmady et al. 2005) |
| RRM1 | CUCUCU | 3.2 μM | 20 mM NaCl | Titration | NMR | (Auweter et al. 2007) |
| RRM2 | CUCUCU | 0.93 μM | 20 mM NaCl | Titration | NMR | (Auweter et al. 2007) |
| RRM34 H457A | CUCUCU | 2.1 μM | 20 mM NaCl | Titration | NMR | (Auweter et al. 2007) |
| RRM34 H411A | CUCUCU | ≤ 2 μM | 20 mM NaCl | Titration | NMR | (Auweter et al. 2007) |
| RRM34 H457A | UCUCU | 23 μM | 20 mM NaCl | Titration | NMR | (Auweter et al. 2007) |
| RRM1 | CUCU | 430 μM | 20 mM NaCl | Titration | NMR | (Auweter et al. 2007) |
| RRM2 | CUCU | 17 μM | 20 mM NaCl | Titration | NMR | (Auweter et al. 2007) |
| RRM34 H411A | CUCU | 46 μM | 20 mM NaCl | Titration | NMR | (Auweter et al. 2007) |
| RRM34 H457A | UUUUU | 110 μM | 20 mM NaCl | Titration | NMR | (Auweter et al. 2007) |
| RRM2 | UUUU | 760 μM | 20 mM NaCl | Titration | NMR | (Auweter et al. 2007) |
| RRM1 | U1 SL4 | 0.85 μM | 20 mM NaCl | Titration | ITC | (Sharma et al. 2011) |
| RRM2 | U1 SL4 | 0.39 μM | 20 mM NaCl | Titration | ITC | (Sharma et al. 2011) |
| RRM1 | UCUCU | 24.7 μM | 20 mM NaCl | Titration | ITC | (Sharma et al. 2011) |
| FL | GABA intron (I) | 5, 130 nM | 100 mM NaCl | 30 mins | EMSA | (Clerte and Hall 2006) |
| RRM12 | GABA intron (I) | 50, 300 nM | 100 mM NaCl | 30 mins | EMSA | (Clerte and Hall 2006) |
| FL | GABA I11/E | 30, 100 nM | 100 mM NaCl | 30 mins | EMSA | (Clerte and Hall 2006) |
| RRM34 | GABA I11/E | 10, 50, 1000 nM | 100 mM NaCl | 30 mins | EMSA | (Clerte and Hall 2006) |
| FL | GABA I34 | 2, 30 nM | 100 mM NaCl | 30 mins | EMSA | (Clerte and Hall 2006) |
| RRM12 | GABA I34 | 50, 160 nM | 100 mM NaCl | 30 mins | EMSA | (Clerte and Hall 2006) |
| RRM34 | GABA I34 | 40, 150 nM | 100 mM NaCl | 30 mins | EMSA | (Clerte and Hall 2006) |
| FL | GABA I11 | 15 | 100 mM NaCl | 30 mins | EMSA | (Clerte and Hall 2006) |
| RRM12 | GABA I11 | Not detected | 100 mM NaCl | 30 mins | EMSA | (Clerte and Hall 2006) |
| RRM34 | GABA I11 | 50 | 100 mM NaCl | 30 mins | EMSA | (Clerte and Hall 2006) |
| PTBP1.4 | CUCAGCCUCUC CUUCUCUCUGC UUCUCUCUCGC UGGCCCUUAG | 302 nM | none | 30 mins | EMSA | (Ontiveros et al. 2020) |
| PTBP1L1Δ40L2Δ70 | CUCAGCCUCUC CUUCUCUCUGC UUCUCUCUCGC UGGCCCUUAG | 24 nM | none | 30 mins | EMSA | (Ontiveros et al. 2020) |
| PTBP1NΔ50L1Δ40L2Δ70 | CUCAGCCUCUC CUUCUCUCUGC UUCUCUCUCGC UGGCCCUUAG | 122 nM | none | 30 mins | EMSA | (Ontiveros et al. 2020) |
| FL | IRES | 1.3 μM | 50 mM NaCl | 10 mins | FDB | (Cobbold et al. 2010) |

| | | | | | | |
|---|---|---|---|---|---|---|
| FL | Mutant IRES | 0.74 μM | 50 mM NaCl | 10 mins | FDB | (Cobbold et al. 2010) |
| FL-Biotinylated | $(CU)_8$ | 38 nM | none | 3 mins | BLI | (Ye et al. 2025) |

**Supplemental Table 4**. TDP43 Construct Binding affinities reported in the literature

| Construct | RNA | $K_D$ | [Salt] | Incubation Time | Method | Citation |
|---|---|---|---|---|---|---|
| RRM1 | $(UG)_3$ | 65 nM | None | Not Reported | FDB | (Kuo et al. 2009) |
| RRM2 | $(UG)_6$ | 379 nM | None | Not Reported | FDB | (Kuo et al. 2009) |
| RRM12 | $(UG)_1$ | N/A | None | Not Reported | FBD | (Kuo et al. 2009) |
| RRM12 | $(UG)_2$ | N/A | None | Not Reported | FBD | (Kuo et al. 2009) |
| RRM12 | $(UG)_3$ | 3060 nM | None | Not Reported | FBD | (Kuo et al. 2009) |
| RRM12 | $(UG)_4$ | 115 nM | None | Not Reported | FBD | (Kuo et al. 2009) |
| RRM12 | $(UG)_6$ | 14.2 nM | None | Not Reported | FBD | (Kuo et al. 2009) |
| RRM12 | $(UG)_8$ | 2.79 nM | None | Not Reported | FBD | (Kuo et al. 2009) |
| RRM12 | CLIP34 | 112 nM | 10 mM NaCl | 20 min | EMSA | (Bhardwaj et al. 2013) |
| RRM12 | CLIP6 | 262 nM | 10 mM NaCl | 20 min | EMSA | (Bhardwaj et al. 2013) |
| RRM12 | CLIP34_UG6 | 5.3 nM | 10 mM NaCl | 20 min | EMSA | (Bhardwaj et al. 2013) |
| FL | $(UG)_5$ | 30 nM | 10 mM NaCl | 15-20 min | EMSA | (Ayala et al. 2005) |
| FL | $(UG)_6$ | 8 nM | 10 mM NaCl | 15-20 min | EMSA | (Ayala et al. 2005) |
| RRM12 | $(GU)_6$ | 22.6 nM | 50 mM KCl | 0-30 min | ITC | (Lukavsky et al. 2013) |
| RRM12 | $A(GU)_6$ | 0.8 nM | 300 mM NaCl | Not Reported | FA | (Grese et al. 2021) |
| NTD-RRM12 | $A(GU)_6$ | 1.1 nM | 300 mM NaCl | Not Reported | FA | (Grese et al. 2021) |
| FL | $A(GU)_6$ | 0.8 nM | 300 mM $Na^+$ | Not Reported | FA | (Grese et al. 2021) |
| RRM12 | AUG12 | 26.9 nM | 50 mM KCl | 0-30 min | ITC | (Lukavsky et al. 2013) |
| FL | $(AU)_{12}$ | 105.3 μM | 50 mM KCl | 20 min | EMSA | (Moore et al. 2026) |
| FL | $(IU)_{12}$ | 2.057 μM, 1.795 μM | 50 mM KCl | 20 min | EMSA | (Moore et al. 2026) |
| FL | $(GU)_{12}$ | 87.5 nM | 50 mM KCl | 20 min | EMSA | (Moore et al. 2026) |
| RRM12 | $(GU)_6$ | 73 pM | 150 mM LiCl | 24 hours | EMSA | (Vivek et al. 2025) |
| RRM12 | $(GU)_6$ | 60 pM | 150 mM NaCl | 24 hours | EMSA | (Vivek et al. 2025) |

| | | | | | | |
|---|---|---|---|---|---|---|
| RRM12 | $(GU)_6$ | 47.9 pM | 150 mM NaCl | 30 minutes | EMSA | (Vivek et al. 2025) |
| RRM12 | $(GU)_6$ - 2AA | 3.9 nM | 150 mM NaCl | 30 minutes | EMSA | (Vivek et al. 2025) |
| RRM12 | $(GU)_6$ - 4AA | N.D. | 150 mM NaCl | 30 minutes | EMSA | (Vivek et al. 2025) |
| RRM12 | $(GU)_6$ - 6AA | 49.4 pM | 150 mM NaCl | 30 minutes | EMSA | https://www.nature.com/articles/s41467-026-76721-y |
| RRM12 | $(GU)_6$ - 8AA | 4.24 nM | 150 mM NaCl | 30 minutes | EMSA | (Vivek et al. 2025) |
| RRM12 | $(GU)_6$ - 10AA | 4.6 nM | 150 mM NaCl | 30 minutes | EMSA | (Vivek et al. 2025) |
| RRM12 | AUG12 | 2.6 nM | 150 mM NaCl | 30 minutes | EMSA | (Vivek et al. 2025) |
| RRM12 | 6 FAM-A | 0.96 nM | 150 mM NaCl | 30 minutes | FP | (Vivek et al. 2025) |
| RRM12 | 6 FAM-A | 0.43 nM | 150 mM NaCl | 30 minutes | FP | (Vivek et al. 2025) |
| RRM12 | 6 FAM-A | 2.8 nM | 150 mM NaCl | 30 minutes | FP | (Vivek et al. 2025) |
| RRM12 | 6 FAM-A | 2.3 nM | 150 mM NaCl | 30 minutes | FP | (Vivek et al. 2025) |
| RRM12 | $(GU)_{12}$ | ≤ 5 pM, 317.8 pM | 150 mM LiCl | 24 hours | EMSA | (Vivek et al. 2025) |
| NTD-RRM12 | $(GU)_6$ | 251 pM | 150 mM LiCl | 12 hours | EMSA | (Vivek et al. 2025) |
| NTD-RRM12 | $(GU)_{12}$ | ≤ 5 pM, 73.7 pM | 150 mM LiCl | 12 hours | EMSA | (Vivek et al. 2025) |
| FL | $(GU)_6$ | 229 pM | 150 mM LiCl | 30 minutes | EMSA | (Vivek et al. 2025) |
| FL-MBP tag | $(GU)_6$ | 1.37 nM | 150 mM LiCl | 30 minutes | EMSA | (Vivek et al. 2025) |
| FL-MBP tag | $(GU)_{12}$ | 88.6, 375 pM | 150 mM LiCl | 30 minutes | EMSA | (Vivek et al. 2025) |

**Supplemental Table 5**. hnRNPU Construct Binding affinities reported in the literature:

| Construct | RNA | $K_D$ | [Salt] | Incubation Time | Method | Citation |
|---|---|---|---|---|---|---|
| U-CTD (673-809) | Syncrip | 82 nM | 150 mM KCl (or LiCl) | 45 mins | FA | (Kletzien et al. 2024) |
| U-CTD | Syncrip G-quad | 80 nM | 150 mM KCl (or LiCl) | 45 mins | FA | (Kletzien et al. 2024) |
| U-CTD | Syncrip GToC | 350 nM | 150 mM KCl (or LiCl) | 45 mins | FA | (Kletzien et al. 2024) |
| U-CTD | Syncrip ΔG-quad | 250 nM | 150 mM KCl (or LiCl) | 45 mins | FA | (Kletzien et al. 2024) |
| U-CTD | SC1 K | 270 nM | 150 mM KCl (or LiCl) | 45 mins | FA | (Kletzien et al. 2024) |
| U-CTD | SC1 Li | 350 nM | 150 mM KCl (or LiCl) | 45 mins | FA | (Kletzien et al. 2024) |
| RGG | Syncrip | 910 nM | 150 mM KCl (or LiCl) | 45 mins | FA | (Kletzien et al. 2024) |
| RGG | Syncrip G-quad | 1600 nM | 150 mM KCl (or LiCl) | 45 mins | FA | (Kletzien et al. 2024) |
| RGG | Syncrip GToC | >10,000 nM | 150 mM KCl (or LiCl) | 45 mins | FA | (Kletzien et al. 2024) |
| RGG | Syncrip ΔG-quad | >10,000 nM | 150 mM KCl (or LiCl) | 45 mins | FA | (Kletzien et al. 2024) |
| RGG | SC1 K | 67 nM | 150 mM KCl (or LiCl) | 45 mins | FA | (Kletzien et al. 2024) |
| RGG | SC1 Li | >19000 nM | 150 mM KCl (or LiCl) | 45 mins | FA | (Kletzien et al. 2024) |
| U-CTD | NEAT1 | 160 nM | 150 mM KCl (or LiCl) | 45 mins | FA | (Kletzien et al. 2024) |
| U-CTD | NEAT1 mutant | 120 nM | 150 mM KCl (or LiCl) | 45 mins | FA | (Kletzien et al. 2024) |
| U-CTD | NEAT1 deletion | 110 nM | 150 mM KCl (or LiCl) | 45 mins | FA | (Kletzien et al. 2024) |
| U-CTD | Antisense NEAT1 | 130 nM | 150 mM KCl (or LiCl) | 45 mins | FA | (Kletzien et al. 2024) |
| U-CTD | MTRNR2L6 | 110 nM | 150 mM KCl (or LiCl) | 45 mins | FA | (Kletzien et al. 2024) |
| U-CTD | MTRNR2L6 mutant | 140 nM | 150 mM KCl (or LiCl) | 45 mins | FA | (Kletzien et al. 2024) |
| U-CTD | MTRNR2L6 deletion | 140 nM | 150 mM KCl (or LiCl) | 45 mins | FA | (Kletzien et al. 2024) |
| U-CTD | Antisense MTRNR2L6 | 170 nM | 150 mM KCl (or LiCl) | 45 mins | FA | (Kletzien et al. 2024) |
| U-CTD | NPM1 | 62 nM | 150 mM KCl (or LiCl) | 45 mins | FA | (Kletzien et al. 2024) |
| U-CTD | NPM1 antisense | 270 nM | 150 mM KCl (or LiCl) | 45 mins | FA | (Kletzien et al. 2024) |
| U-CTD | KCNQ1OT1 | 94 nM | 150 mM KCl (or | 45 mins | FA | (Kletzien et al. 2024) |

| | | | | | | |
|---|---|---|---|---|---|---|
| | | | LiCl) | | | |
| U-CTD | KCNQ1OT1 antisense | 160 nM | 150 mM KCl (or LiCl) | 45 mins | FA | (Kletzien et al. 2024) |
| U-CTD | hnRNPA1 | 150 nM | 150 mM KCl (or LiCl) | 45 mins | FA | (Kletzien et al. 2024) |
| U-CTD | hnRNPA1 antisense | 140 nM | 150 mM KCl (or LiCl) | 45 mins | FA | (Kletzien et al. 2024) |
| U-CTD | KCNQ1 | 960 nM | 150 mM KCl (or LiCl) | 45 mins | FA | (Kletzien et al. 2024) |
| U-CTD | KCNQ1 antisense | 1800 nM | 150 mM KCl (or LiCl) | 45 mins | FA | (Kletzien et al. 2024) |
| U-CTD | Syncrip antisense | 280 nM | 150 mM KCl (or LiCl) | 45 mins | FA | (Kletzien et al. 2024) |
| U(673-825) | Syncrip | 82 nM | 150 mM KCl (or LiCl) | 45 mins | FA | (Kletzien et al. 2024) |
| U(702-825) | Syncrip | 70 nM | 150 mM KCl (or LiCl) | 45 mins | FA | (Kletzien et al. 2024) |
| U(702-809) | Syncrip | 140 nM | 150 mM KCl (or LiCl) | 45 mins | FA | (Kletzien et al. 2024) |
| U(702-793) | Syncrip | 180 nM | 150 mM KCl (or LiCl) | 45 mins | FA | (Kletzien et al. 2024) |
| U(702-777) | Syncrip | 450 nM | 150 mM KCl (or LiCl) | 45 mins | FA | (Kletzien et al. 2024) |
| U(702-761) | Syncrip | 1100 nM | 150 mM KCl (or LiCl) | 45 mins | FA | (Kletzien et al. 2024) |
| U(702-745) | Syncrip | 910 nM | 150 mM KCl (or LiCl) | 45 mins | | (Kletzien et al. 2024) |
| U(673-825) | Syncrip antisense | 280 nM | 150 mM KCl (or LiCl) | 45 mins | FA | (Kletzien et al. 2024) |
| U(702-825) | Syncrip antisense | 570 nM | 150 mM KCl (or LiCl) | 45 mins | FA | (Kletzien et al. 2024) |
| U(702-809) | Syncrip antisense | 1600 nM | 150 mM KCl (or LiCl) | 45 mins | FA | (Kletzien et al. 2024) |
| U(702-793) | Syncrip antisense | 3000 nM | 150 mM KCl (or LiCl) | 45 mins | FA | (Kletzien et al. 2024) |
| U(702-777) | Syncrip antisense | 14000 nM | 150 mM KCl (or LiCl) | 45 mins | FA | (Kletzien et al. 2024) |
| U(702-761) | Syncrip antisense | >21000 nM | 150 mM KCl (or LiCl) | 45 mins | FA | (Kletzien et al. 2024) |
| U(702-745) | Syncrip antisense | >26,000 nM | 150 mM KCl (or LiCl) | 45 mins | FA | (Kletzien et al. 2024) |
| U(673-825) | MTRNR2L6 | 110 nM | 150 mM KCl (or LiCl) | 45 mins | FA | (Kletzien et al. 2024) |
| U(702-825) | MTRNR2L6 | 220 nM | 150 mM KCl (or LiCl) | 45 mins | FA | (Kletzien et al. 2024) |
| U(702-809) | MTRNR2L6 | 710 nM | 150 mM KCl (or LiCl) | 45 mins | FA | (Kletzien et al. 2024) |

| | | | | | | |
|---|---|---|---|---|---|---|
| U(702-793) | MTRNR2L6 | 1100 nM | 150 mM KCl (or LiCl) | 45 mins | FA | (Kletzien et al. 2024) |
| U(702-777) | MTRNR2L6 | 4200 nM | 150 mM KCl (or LiCl) | 45 mins | FA | (Kletzien et al. 2024) |
| U(702-761) | MTRNR2L6 | > 11,000 nM | 150 mM KCl (or LiCl) | 45 mins | FA | (Kletzien et al. 2024) |
| U(702-745) | MTRNR2L6 | >13,000 nM | 150 mM KCl (or LiCl) | 45 mins | FA | (Kletzien et al. 2024) |
| U(673-825) | Antisense MTRNR2L6 | 160 nM | 150 mM KCl (or LiCl) | 45 mins | FA | (Kletzien et al. 2024) |
| U(702-825) | Antisense MTRNR2L6 | 450 nM | 150 mM KCl (or LiCl) | 45 mins | FA | (Kletzien et al. 2024) |
| U(702-809) | Antisense MTRNR2L6 | 810 nM | 150 mM KCl (or LiCl) | 45 mins | FA | (Kletzien et al. 2024) |
| U(702-793) | Antisense MTRNR2L6 | 1200 nM | 150 mM KCl (or LiCl) | 45 mins | FA | (Kletzien et al. 2024) |
| U(702-777) | Antisense MTRNR2L6 | 3900 nM | 150 mM KCl (or LiCl) | 45 mins | FA | (Kletzien et al. 2024) |
| U(702-761) | Antisense MTRNR2L6 | >9,000 nM | 150 mM KCl (or LiCl) | 45 mins | FA | (Kletzien et al. 2024) |
| U(702-745) | Antisense MTRNR2L6 | >13,000 nM | 150 mM KCl (or LiCl) | 45 mins | FA | (Kletzien et al. 2024) |
